# A multiscale stratigraphic investigation of the context of StW 573 ‘Little Foot’ and Member 2, Sterkfontein Caves, South Africa

**DOI:** 10.1101/482711

**Authors:** Laurent Bruxelles, Dominic J. Stratford, Richard Maire, Travis R. Pickering, Jason L. Heaton, Amelie Beaudet, Kathleen Kuman, Robin Crompton, Kris J. Carlson, Tea Jashashvili, Juliet McClymont, George M. Leader, Ronald J. Clarke

## Abstract

The Sterkfontein Caves has an 80 year history of yielding remarkable evidence of hominin evolution and is currently the world’s richest *Australopithecus-beafing* site. Included in Sterkfontein’s hominin assemblage is StW 573 (‘Little Foot’). Discovered in the Member 2 deposit in the Silberberg Grotto, StW 573 represents the most complete *Australopithecus* skeleton yet found. Because of its importance to the fossil hominin record, the geological age of Little Foot has been the subject of significant debate. Two main hypotheses have been proposed regarding the formation and age of Member 2 and by association StW 573. The first, proposes that Member 2 formed relatively rapidly, starting to accumulate at around 2.8 million years ago and that the unit is isolated to the Silberberg Grotto - the underlying chambers and passages forming later. The second proposes that Member 2 formed slowly, its accumulation starting before 3.67 million years ago and that the deposit extends into the Milner Hall and close to the base of the cave system. Both assume a primary association between StW 573 and Member 2, although which sediments in the Silberberg Grotto are associated with Member 2 has also been questioned. Recently a third alternative hypothesis questioning the association of StW 573 to Member 2 sediments proposed a ‘two-stage burial scenario’ in which sediments associated with StW 573 represent a secondary and mixed-age deposit reworked from a higher cave. The stratigraphic and sedimentological implications of these hypotheses are tested here through the application of a multiscale investigation of Member 2, with reference to the taphonomy of the Little Foot skeleton. The complete infilling sequence of Member 2 is described and depositional units are tracked across all exposures of the deposit in the Silberberg Grotto and into the Milner Hall. Facies development follows patterns characteristic of colluvially accumulated taluses with 30-40° angles of repose developing coarser distal facies. Sediments are generally stratified and conformably deposited in a sequence of silty sands eroded from well-developed lateritic soils on the landscape surface. Voids, clasts and bioclasts are organized consistently across and through Member 2 according to the underlying deposit geometry, indicating a gradual deposit accretion with no distinct collapse facies evident, no successive debris flow accumulation, and only localized intra-unit post-depositional modification. The stratigraphy and sedimentology of Member 2 supports a simple single-stage accumulation process through which Member 2 partially fills the Silberberg Grotto and extends into the deeper chambers and passages of the Sterkfontein Caves. Through this work we demonstrate at multiple scales the primary association between the sediments of Member 2 and the StW 573 ‘Little Foot’ skeleton.

## 1. Introduction

The Sterkfontein Caves, located about 50km north-west of Johannesburg, South Africa (Stratford & Crompton, 2018) are situated on the southern side of the Bloubank River in the south-western area of the Cradle of Humankind. Over the last 80 years, the caves of the Cradle of Humankind (COH) have yielded some of the most significant hominin specimens representing multiple species and attributed to three genera of hominin. Research at Sterkfontein has yielded specimens representing three genera of hominin and potentially five species spanning possibly 4 million years (maximum age based on Partridge et al., 2003, see below for discussion). The depositional sequence used by researchers to divide the Sterkfontein sequence into chronostratigraphic units was developed by Partridge (1978; 2000) and is referred to as the Sterkfontein Formation (also see Stratford, 2017 for review). This sequence comprises six members ordered stratigraphically (although in no single exposure are all members represented), with Member 1 at the base and Member 6 at the top. Members 2 through 6 represent ‘allogenic’ fillings derived from outside the cave. Member 1, however, consists of an ‘autogenic’ filling developed internally through cave breakdown before an opening to the surface had formed. The majority of hominin specimens found at Sterkfontein are attributed to the *Australopithecus* genus and were recovered from the Member 4 deposits of the Sterkfontein Formation (Partridge, 1978). Deeper chambers have yielded *Australopithecus* specimens of potentially Pliocene age (Clarke, 1998; Partridge et al., 2003; see below for discussion of geological ages of specimens). The most complete of these deeper specimens is StW 573 (‘Little Foot’), a near-complete skeleton dated to 3.67 million years old (Granger et al., 2015), excavated from the Member 2 deposit found in the Silberberg Grotto (Clarke, 1998). For a map of the Sterkfontein Cave system, see Stratford & Crompton (2018, in this issue).

Sterkfontein Member 2 has been the subject of intense debate since the discovery of the StW 573 ‘Little Foot’ (Clarke, 1998). Debate has centered largely around the age of the specimen (Clarke & Tobias, 1995; McKee, 1996; Tobias & Clarke, 1996; Turner, 1997; Partridge et al., 1999, 2003; Berger et al., 2002; Clarke, 2002; Walker et al., 2006; Pickering & Kramers, 2010; Herries & Shaw, 2011; Granger et al., 2015; Kramers & Dirks, 2017a,b; Stratford et al., 2017), and it has necessarily included discussion of the stratigraphic history of StW 573 and Member 2 (Clarke, 1998, 2002a,b, 2006, 2007, 2008; Bruxelles et al., 2014;

Pickering & Kramers, 2010; Bruxelles et al., 2014; Kramers & Dirks, 2017a,b; Stratford et al., 2017). Member 2 is the basal allogenic unit of the Sterkfontein Formation. It was originally described by Partridge (1978) from exposures in the Silberberg Grotto as one of the deeper chambers in the Sterkfontein cave system but overlying five km of passages and chambers (including the Milner Hall and Elephant Chamber) (Martini et al., 2003). Although debate continues about the geological age of Member 2, some data suggest it may represent the oldest *Australopithecus-bearing* deposit included in the Sterkfontein Formation (e.g., Clarke, 2002; Partridge et al., 2003; Granger et al., 2015). The geological age of Member 2 and StW 573 has significant implications for not only where Little Foot lies in hominin phylogeny, but also for the maximum age of deposits at Sterkfontein and the process of formation of the caves themselves. Various hypotheses have been proposed regarding the geological age of StW 573 and its stratigraphic history in relation to the evolution of the caves. These are summarized here and discussed in detail below. Pickering and Kramers (2010) proposed: that Member 2 accumulated rapidly from the surface from about 2.8 million years ago; that some of the sediments in the western Silberberg Grotto may not be Member 2; and that Member 2 is restricted to the Silberberg Grotto, with the lower chambers and passages forming after the Silberberg Grotto. This final point supports proposals by Partridge (1978) and Partridge and Watt (1991) for an epiphreatic karstification process forming the caves, i e., deeper networks are formed later and necessarily filled with younger or reworked sediments.

Granger et al. (2015), applied cosmogenic nuclide dating to nine samples comprising clasts and sediments stratigraphically associated with StW 573. The results of the analysis supported a geological age of 3.67 Ma for the sediments containing Little Foot. From the cosmogenic samples, Granger et al. (2015) proposed a slow landscape erosion rate, and therefore slow rate of Member 2 accumulation. Kramers and Dirks (2017a,b) suggested that one of the nine samples included in the cosmogenic nuclide isochron published by Granger et al. (2015) could indicate re-working of some sediments in Member 2. They argued that a two-stage burial scenario involving a collapse of sediments from an upper chamber containing the Little Foot skeleton would explain this one anomalous sample (see Stratford et al., 2017 for reply).

Sedimentological evidence from exposures of Member 2 in the Silberberg Grotto deposit should preserve evidence of the mode of accumulation of the deposit and can therefore be used to test the above hypotheses. Refining the stratigraphic history of Member 2 improves our understanding of the formation and opening of the caves and the local geomorphological, ecological and environmental conditions during the earliest infilling, thereby providing a more nuanced contextual framework for hominin evolution in southern Africa during the Plio-Pleistocene. In addition, understanding the process of accumulation of Member 2 and the associated taphonomic implications for the Little Foot skeleton, we can test the hypotheses presented by Pickering and Kramers (2010) and Kramers and Dirks (2017a,b) and associate sedimentological features to depositional processes and surface geomorphological contexts. Research presented here draws on multiscale and multidisciplinary evidence to further clarify the stratigraphic context of Member 2 and the association of StW 573 with that unit. It is our goal to integrate these data to provide a more holistic perspective on deposit development that can be correlated with ongoing broader landscape evolution research in the southern Cradle of Humankind landscape.

## 2. Background: Member 2 Stratigraphy, Age, Taphonomy and Palaeoenvironments

### 2.1 Previous interpretations of Member 2 stratigraphy

Partridge (1978) first described the Member 2 deposit from exposures in the Silberberg Grotto (Figure 1; and Stratford & Crompton, 2018). The morphology of the Silberberg Grotto is defined by the strong east to west fault-guided cavity development that characterizes the Sterkfontein system (Martini et al., 2003; Stratford, 2017; Bruxelles 2018). The chamber has developed along the southern boundary of the cave system and extends approximately 30 m east to west. To the west, the chamber divides into three smaller passages that continue west and open into the underlying Elephant Chamber and Milner Hall (Figure 1). The northern and eastern extents of the chamber are unknown as these walls are comprised of breccia^1^ and mined speleothem remnants, but based on the proximity of the Name Chamber to the north (Martini et al., 2003) it is likely the chamber’s northern extent is limited to less than 20 m from the southern wall. The large talus cone of Member 2 was deposited on the thick flowstone covering the irregular surface of the autogenic Member 1 (Partridge, 1978, 2000; Martini et al., 2003). Several researchers have described the morphology of Member 2 in relation to its formation and proposed that the deposit accumulated from an aven-like opening high in the roof above the eastern area of the chamber (Partridge, 1978; Martini et al., 2003; Clarke, 2006). The east and west flanks of Member 2 slope away from this point (Figure 1). StW 573 was found about 18 m down the steeper western slope (Clarke, 2006; Bruxelles et al., 2014) (Figure 1).

**Figure 1:**
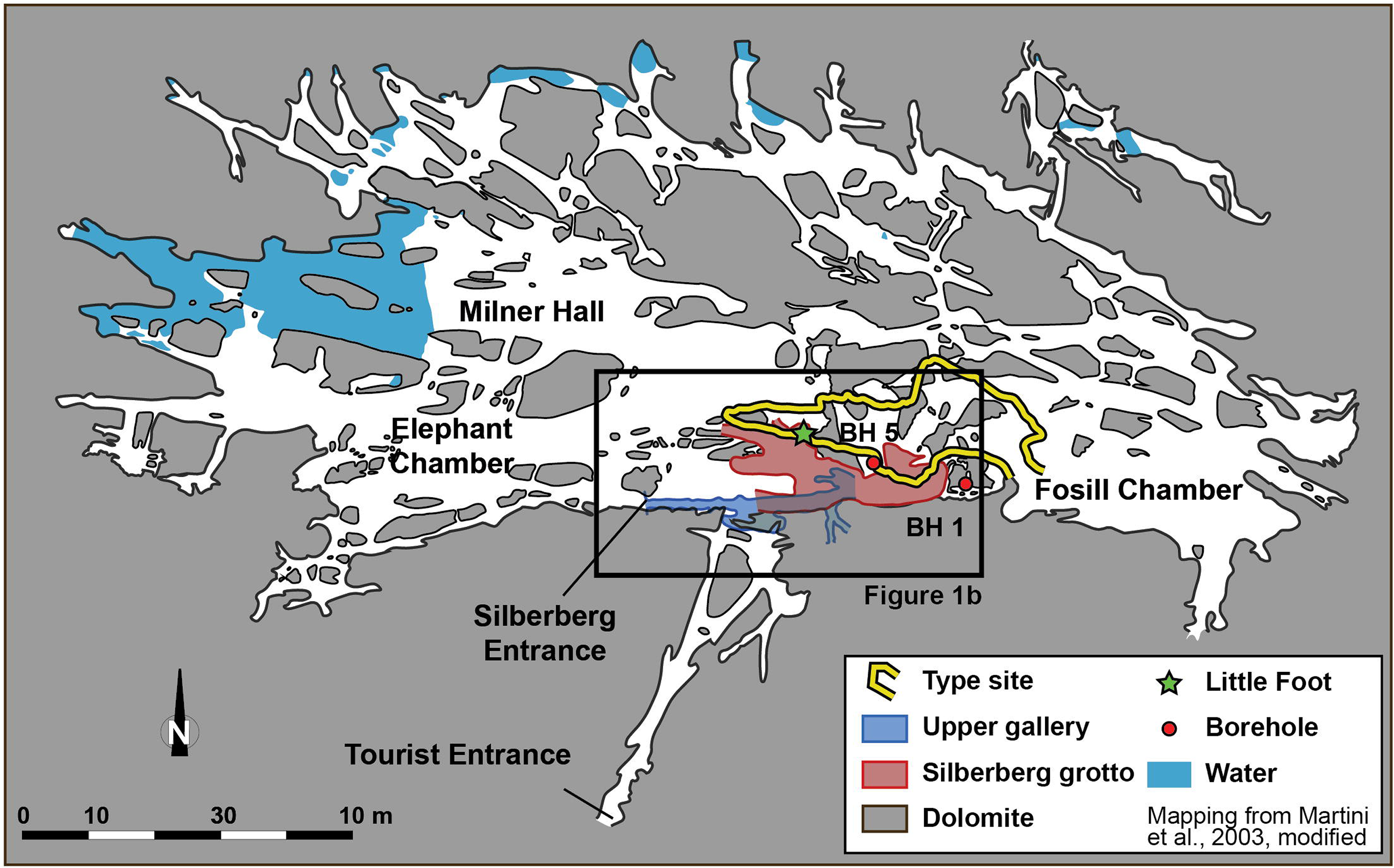
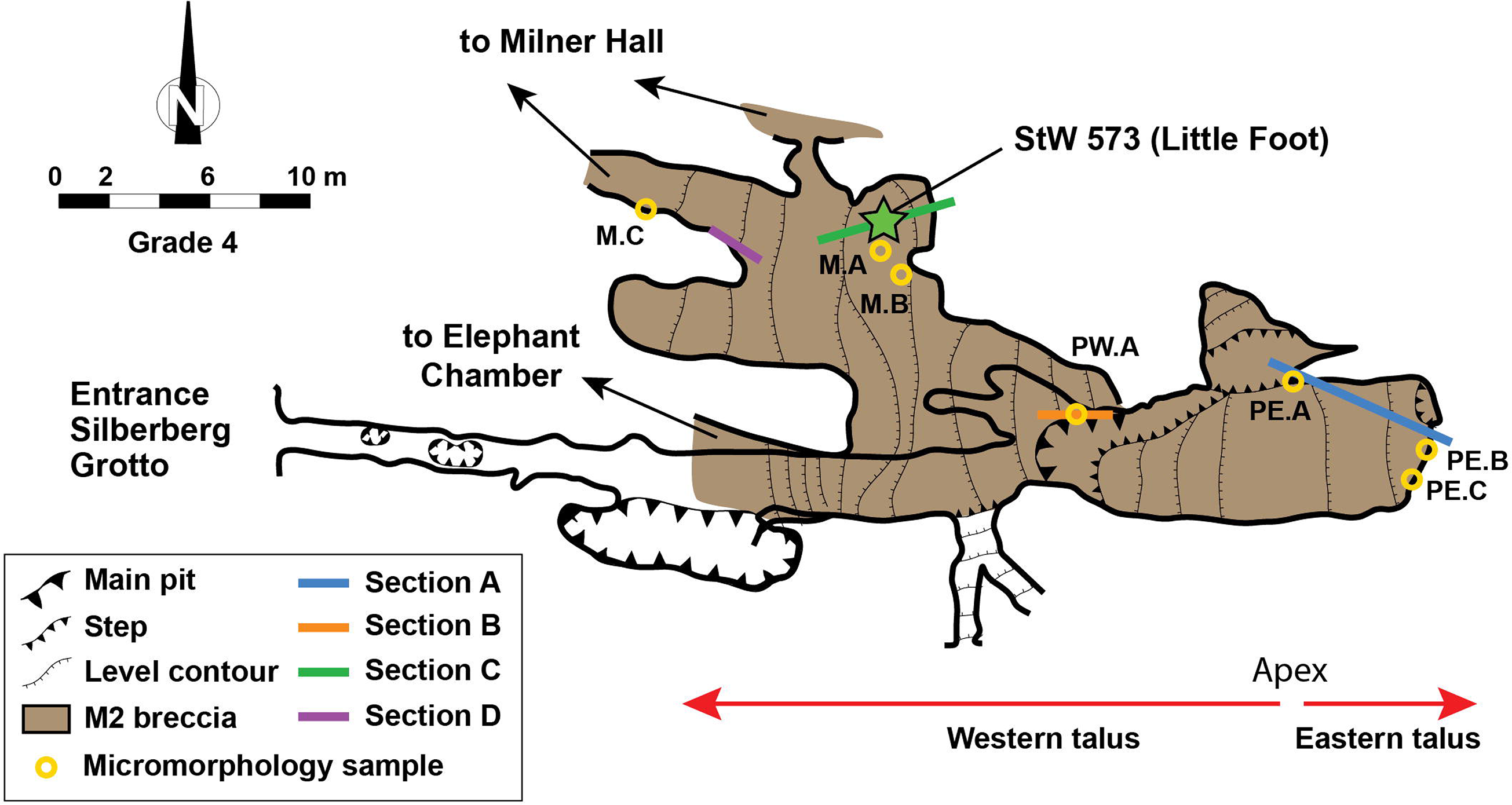
Plan of the Silberberg Grotto with openings to associated chambers, talus topography, locations of described sections and sediment micromorphology samples. Star indicates location of StW 573. Adjusted from Bruxelles et al. (2014).

In his seminal stratigraphic work, Partridge (1978) placed Member 2 at the base of the fossiliferous sequence. Underlying Member 2 is Member 1 and a variably associated speleothm. This unit has been most notably described, albeit briefly, by Partridge (1978; 2000), Partridge and Watt (1991), Martini (2003), Clarke (2006) and Pickering and Kramers (2010). The clastic component of Member 1 is consistently described as a collapse breccia dominated by unorganized, angular dolomitic and chert blocks in a manganiferous matrix containing stalagmitic and lenticular speleothems. The unit was formed through the vadose breakdown of the cave prior to significant opening to the surface. Partridge (1978) originally recognized a contamination of allogenic clastic material and bone in the upper reaches, also identified in cores by Pickering and Kramers (2010), but this is not noted in Martini (2003) or Clarke (2006). The contact between Member 1 and the overlying Member 2 is variable. In boreholes 1 and 4 (Partridge and Watt, 1991; Pickering and Kramers, 2010), a speleothem unit evidently separates Member 1 from 2, but from exposures in the Silberberg Grotto, Partridge (1978) does not describe a speleothem present at the contact, although he does describe “a large stalagmitic boss belonging to Member 1” (pp. 284), and Clarke (2006) describes Member 1 as being ‘cemented with flowstone’ (pp. 115). Clarke proposes the general formation of a speleothem ‘boss’ on Member 1, on which Member 2 formed. Where this speleothem is present and interstratifying Members 1 and 2, we associate this speleothem with a stage of Member 1 formation during a period prior to significant opening of the caves to the surface. Hence this is a pre-Member 2 unit. We refer to this speleothem as a ‘boss’ speleothem.

From exposures in the eastern Silberberg Grotto and the ‘east pit of the type site’, Partridge (1978, p. 284) identified a thick allogenic unit overlying Member 2, which he named Member 3. A flowstone unit named 3A separated the two members (Partridge, 1978; Partridge and Watt, 1991) and was said to be represented in boreholes 1, 4 and perhaps 5 (Pickering and Kramers, 2010). Martini et al. (2003) describe this unit as a relatively regular and widespread flowstone sheet. Member 3 is not visible in the western Silberberg Grotto, and the interbedding flowstone named by Partridge as Unit 3A is also absent in the western part of the grotto. We, therefore, do not discuss these deposits in the work presented here. However, the stratigraphic sections we present document the contact between Member 2 and Member 3. While Pickering and Kramers (2010) have questioned Member 3 as being distinct from the overlying Member 4, we do not attempt to address this question as our focus is on Member 2.

Member 2 was described as a well-bedded, silty-loam matrix-supported talus of up to 5 m depth, with ‘sparse rock debris’ and localized abundant bone. The western slope was interpreted by Partridge (1978, p. 284) as accumulating through a “conical gravitative accretion.” On the eastern side of the talus apex (Figure 1), the bone-rich shallower dipping eastern flank was deposited under fluid-driven sedimentation (Partridge, 1978; Partridge and Watt, 1991). Partridge (1978, 2000) clearly identifies the upper limit of Member 2 in the eastern Silberberg Grotto as sealed by a flowstone that represents the first unit of Member 3 (‘Bed A’ in Partridge (1978), ‘SA-7’ in our sections). The clastic sediments of Member 2 and Member 3 are distinctly separated in the Silberberg Grotto, and we consider them different deposits. Below we summarize work referring to Member 2 and do not consider Member 3 as playing a role in the mode of deposition or morphology of the stratigraphically lower Member 2.

Following the discovery of StW 573 Clarke described the specimen as associated with a stony breccia with layers of reddish, dark brown mudflows, calcite flowstones and underlying cavities (Clarke, 1998, 2002b, 2006), proposing a detailed formation scenario for Member 2 and the sediments directly associated with StW 573. Martini et al. (2003) estimated the maximum thickness of Member 2 as 8 m and described the deposit similarly to Partridge (1978) and Partridge and Watt (1991), but he placed greater emphasis on deposition of sediment by “stream action” (Martini et al., 2003; pp. 58). Pickering and Kramers (2010) also proposed a possible maximum thickness of the Member 2 of 8 m (in Borehole 1) and describe the deposit as a “Pocket of coarse-grained, blocky dolomite-rich rock debris in a reddish-brown sandy matrix with several inter-bedded flowstone layers” (Table 6; p. 81). However, they also noted that Borehole 1 does not transect the Silberberg Grotto and so may not reach or sample what was considered to be Member 2 in the original description.

Pickering and Kramers (2010) attribute the sediments around StW 573 to ‘Facies A’ - interpreted as a proximal talus facies near the StW 573 specimen. More eastern sediments were attributed to ‘Facies C’ - interpreted as distal talus facies. This suggests that the sediments containing StW 573 were associated with a local ‘pocket’ of proximal Member 2 sediments closely associated with distal sediments of another deposit (‘Member *2?*) (see Fig. 3 of Pickering & Kramers, 2010, p. 74).

**Figure 2:**
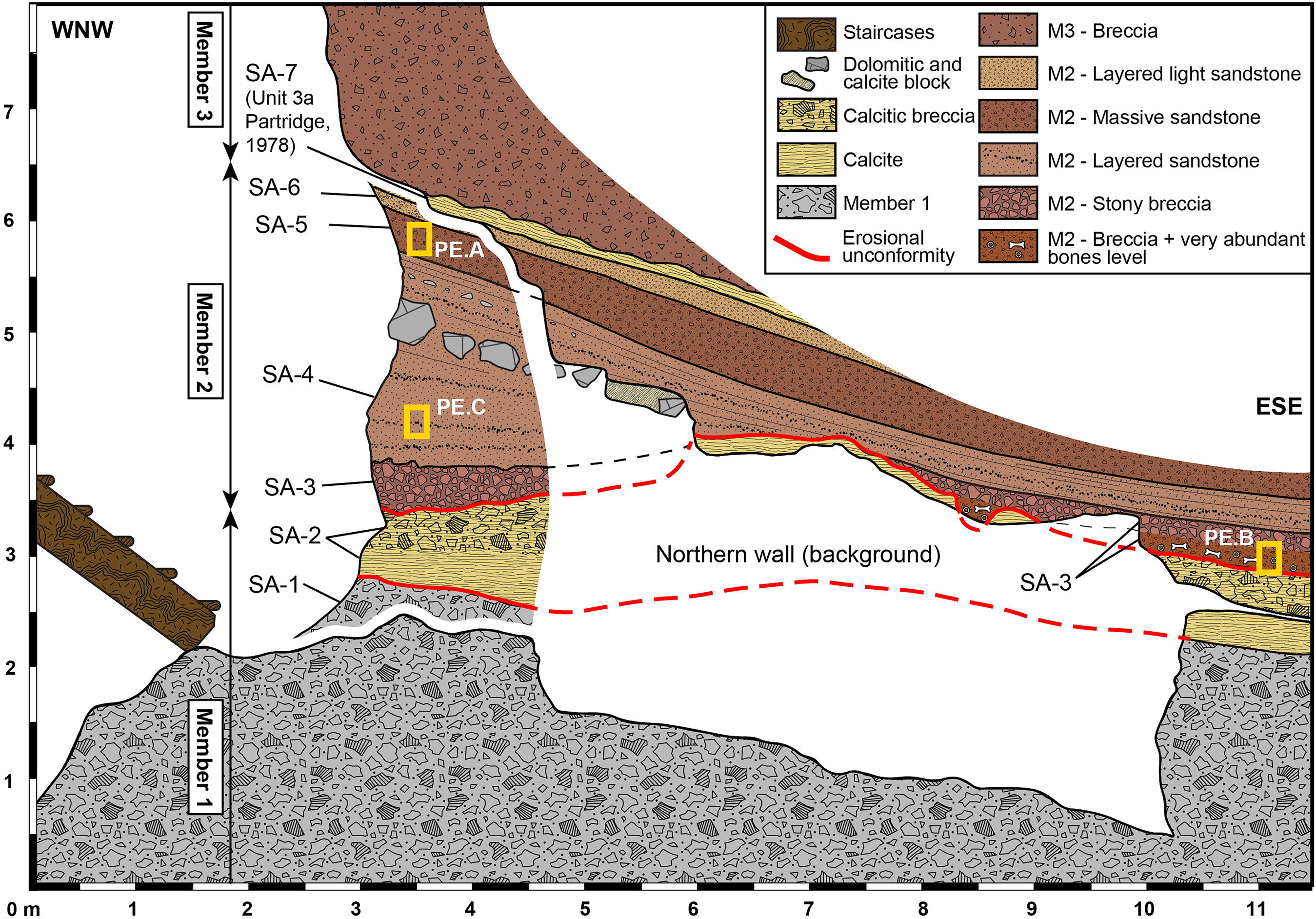
Synthetic representation of Section A (see Figure 1 for location).

**Figure 3:**
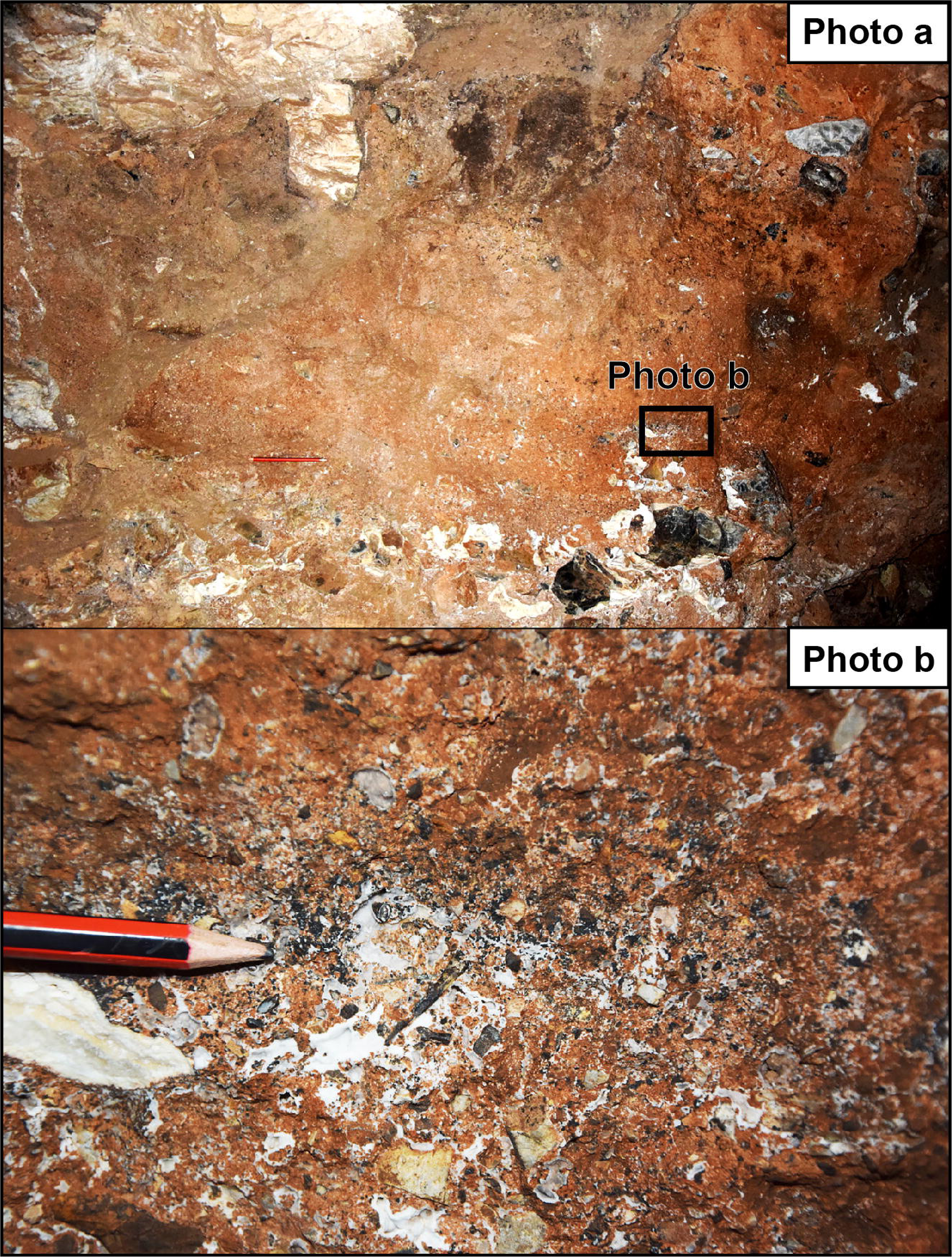
Section A of the Eastern talus (photo a): Stratified deposits with microfauna rich layers (photo b).

**Figure 4:**
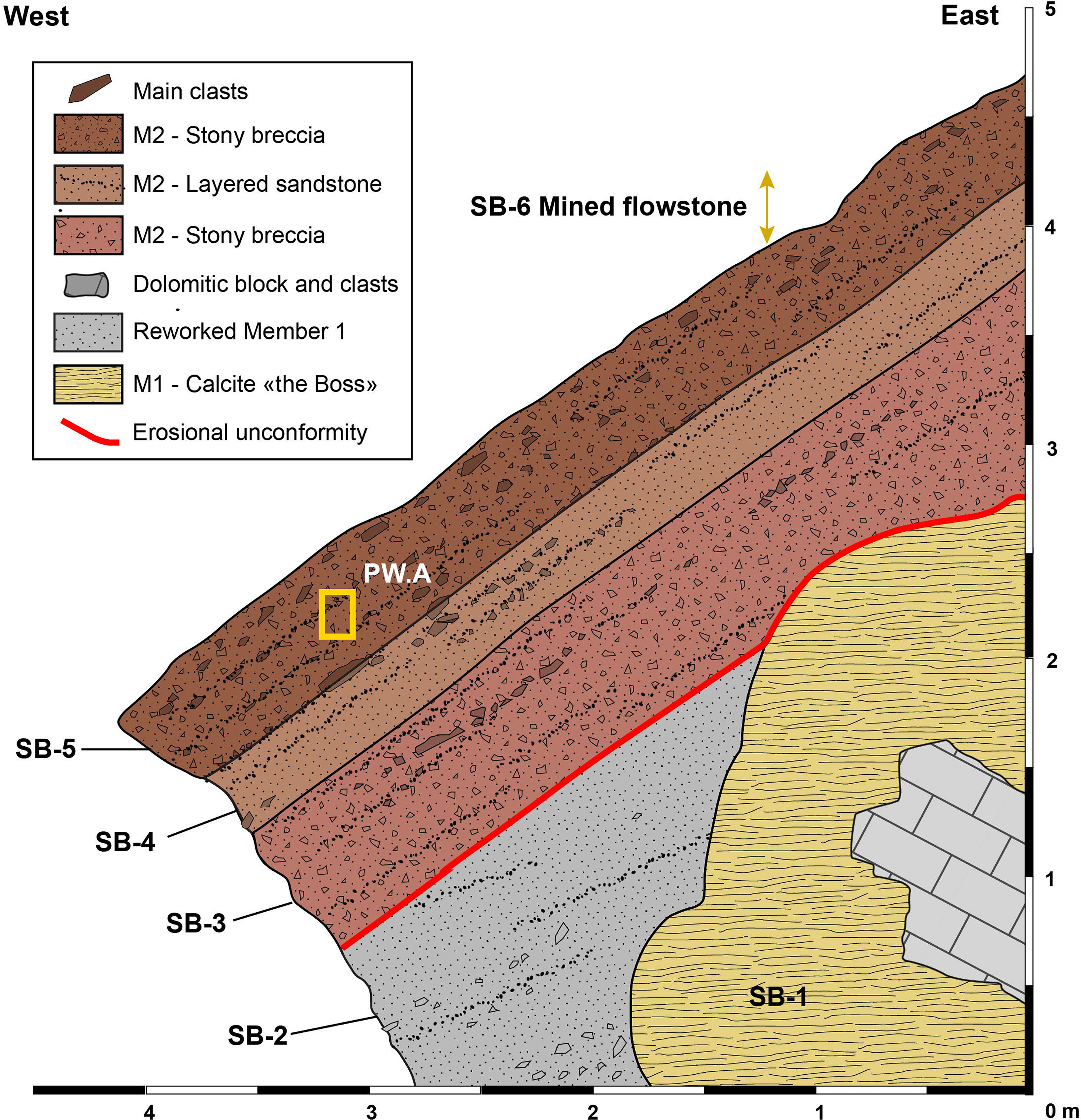
Synthetic representation of Section B (see Figure 1 for location).

In contrast, Bruxelles et al. (2014) examined the sediments of Member 2 associated with the StW 573 skeleton and identified a stratified sequence of three clastic units (B1, B2a,b & B3) and five speleothemic units (F1, F2, F3, F4a,b & F5). Clasts increase in abundance in B2b and B3, but the matrix is generally identified as ‘clayey sand’. Localized cavities have formed in all clastic units which were subsequently filled with speleothem, a process observed by Clarke (2002b) and documented in detail by Bruxelles et al. (2014). Such cavities are linked to episodic increases in fresh water throughput, localized pooling of water, and flushing of poorly indurated sediments (Partridge, 1978; Clarke, 2002b, 2006; Martini et al., 2003; Bruxelles et al., 2014). These processes caused the localized vertical displacement of the central part of the StW 573 skeleton.

Regarding the possible extension of Member 2 beyond the Silberberg Grotto, Partridge (1978) and Partridge and Watt (1991) considered the underlying passages and chambers, like the Elephant Chamber and Milner Hall to have formed after the Silberberg Grotto through a lowering of the base level associated with an epiphreatic karstification model. They thus considered that all deposits contained in the chambers and passages underlying the Silberberg Grotto must be either formed from reworked old sediments, or younger sediments deposited to the base of the system through the numerous aven-style openings that currently feed sediments from the surface to the lower levels of the cave system. This theory was supported by U-Pb dates of younger than 400 ka for some flowstones covering talus deposits found near the base of the cave system in the Milner Hall and Jacovec Cavern (Pickering & Kramers, 2010).

Wilkinson’s geomorphological work (1973, 1983, 1985) argued that the Jacovec Cavern sediments and Silberberg Grotto deposits extend close to the current base level. He thus implied that the deepest deposits, which are found in the Jacovec Cavern and Milner Hall (as an extension of the Member 2 deposit exiting the Silberberg Grotto), are some of the oldest. This model has been supported by recent stratigraphic work conducted in the Milner Hall (Stratford et al., 2014) and by preliminary cosmogenic nuclide dates of 3.76 - 4.02 Ma for hominin-bearing deposits in the Jacovec Cavern (Partridge et al., 2003). This has important implications for the potential extent (depth and distribution) of Member 2, which should be considered when assessing the relationship between the age and formation hypotheses of the deposit and caves.

### 2.2. Member 2 age and implications for deposit formation

Significant attention has centered on the dating of the StW 573 skeleton. Application of a range of methods including biochronology (e.g. Clarke & Tobias, 1995; McKee, 1996; Turner, 1997; Clarke, 1998; Berger et al., 2002; Clarke, 2002a), U-series dating (e.g. Walker et al., 2006; Pickering & Kramers, 2010), palaeomagnetic dating (e.g. Partridge et al., 1999, 2000; Herries and Shaw, 2011) and cosmogenic nuclide dating (Partridge et al., 2003; Granger et al., 2015) has produced dates from a maximum of 4 Ma (Partridge et al., 2003) to a minimum of 1.07 Ma (McKee, 1996; Berger et al., 2002; see Clarke & Tobias, 1996 and Clarke, 2002a for respective replies). Palaeomagnetic dating has yielded various dates from 3.33 Ma (Partridge et al., 1999) to 2.2 Ma, with a proposed date for Little Foot of no older than 2.58 Ma (Herries and Shaw, 2011). U-series and palaeomagnetic work based on samples from flowstones closely associated with StW 573 have generally yielded dates of between 2.8 and 2.6 to 2.16 Ma, with flowstones in the Silberberg Grotto producing U-Pb dates of 2.2 Ma (Walker et al., 2006; Pickering and Kramers, 2010), as opposed to the 2.8 Ma flowstones identified in Boreholes 1 and 4, which are proposed to correlate with an upper limit of Member 2 but, as mentioned above, the boreholes do not transect the Silberberg Grotto (Pickering and Kramers, 2010; Herries and Shaw, 2011; see Herries et al., 2013 for synopsis). Although authors acknowledge that the 2.2 Ma flowstone formed after the deposition of the skeleton, the simple bounding stratigraphic association of these flowstones as suitable for dating the StW 573-bearing sediments was challenged (Clarke, 2002b, 2006; Bruxelles et al., 2014). In other words, stratigraphic considerations of these flowstones presented by Clarke (2002b) and Bruxelles et al. (2014) demonstrated that they are post-depositional void infills that cannot directly date the skeleton. Recent application of a refined cosmogenic nuclide dating method yielded a date of 3.67 Ma for the deposition of the sediments and clasts stratigraphically associated with the specimen (Granger et al., 2015). The association of some of these samples with the skeleton was recently questioned by Kramers and Dirks (2017a,b), who suggested that the older sediments were reworked through the collapse of an upper chamber which also carried with them a younger StW 573 (see Stratford et al., 2017 for reply). The result of these debates has been the emergence of two stances, a younger age of between 2.8 and 2.2 Ma and an older age in excess of 3.510 Ma (based on the younger age within the 0.16 Ma error range of Granger et al,. 2015). The recent proposal of Kramers and Dirks (2017a,b) provides an additional formation hypothesis specifically for those sediments close to the StW 573 specimen and must also be considered.

This article utilizes sedimentological and stratigraphic evidence at multiple scales to clarify the formation processes of Member 2 and relate those processes to the association and taphonomic history of the StW 573 skeleton. The stratigraphic control will help relate the formation processes to the greater system evolution and help test the physical manifestations of the above dating hypotheses (e.g., evidence of localized collapse, rapidity of sedimentation, landscape stability, and sediment provenience).

### 2.3. Member 2 taphonomy

Pickering et al. (2004) identify the presence of antimeric and articulating specimens and the absence of carnivore modification, juvenile (dependent age) carnivore specimens, coprolites, and digested bone as evidence of an assemblage primarily accumulated through ‘death-trap’ processes. Extinct genera of primates (*Parapapio*), carnivores *(Chasmaporthetes)* and bovids (*Makapania)* are present in an assemblage overwhelmingly dominated by primates and felids (Pickering et al., 2004). Pickering et al. (2004) suggest that animals with ‘climbing proclivities’ entered the Silberberg Grotto on their own, either by falling in or by entering an entrance high in the ceiling of the Silberberg Grotto and subsequently being unable to escape. Very limited carnivore involvement is demonstrated by the very low percentages of such modifications—less than 6% of the primate assemblage and less than 3% of the non-primate assemblage (Pickering et al., 2004). There is no evidence that points to a substantial role for carnivores in the accumulation process (Pickering et al., 2004). The presence of a complete and articulated *Australopithecus* skeleton with no observable biogenic modification (Pickering et al., 2004) supports the argument for a mainly death-trap accumulation process. In contrast with Member 4, StW 573 is the only hominin represented in the current Member 2 assemblage. Together, the skeleton from the western talus and the articulating remains of fauna from Dump 20 and from in situ blasting of the eastern talus show a consistent accumulation mode across the Member 2 deposit (Pickering et al., 2004; Clarke, 2006).

### 2.4. Member 2 palaeoenvironments

The cercopithecoid fauna of Member 2 is dominated by small-bodied papionins, such as *Parapapio jonesi, Parapapio broomi* and *Papio izodi,* along with one colobine species, *Cercopithecoides williamsi* (Heaton, 2006; Pickering et al., 2004). The Sterkfontein papionins were much smaller than modern-day species and exhibited lesser degrees of sexual dimorphism (Heaton, 2006, 2007). In contrast, fossil colobines were significantly larger (Delson et al., 2000) and have been used to argue for a clear terrestrial component in assemblages from eastern Africa (Jablonski et al., 2008). While *Cercopithecoides* from southern Africa are smaller, conclusions about their terrestrial proclivities have been extended to sites within the Sterkfontein valley (Ciochon, 1993; Elton, 2001). However, data from the papionins appear mixed. The post-cranial anatomy of *Parapapio jonesi* has been argued to indicate open areas, while the slightly larger *Parapapio broomi* may have been arboreal (Elton, 2001). However, Maier (1970) concluded that *Cercopithecoides williamsi* would have lived near densely forested areas, possibly a tropical fringe forest along rivers of the nearby valleys. Similarly, Elton et al. (2016) argued that the species of Sterkfontein Member 2 were probably to some extent ecologically dependent upon trees for foraging and predator avoidance, or both. Among the papionins, a shift to larger population body size, and perhaps greater territoriality, is not strongly indicated until Member 5 of the Sterkfontein Formation when there was a drier climate (Luyt & Lee-Thorp, 2003; Heaton, 2006, 2007).

The Member 2 mammalian fauna includes caracals, *Makapania,* and monkeys and indicates a palaeohabitat of rocky hills covered in brush and scrub, but valley bottoms with riverine forest, swamp and standing water (Pickering et al., 2004). The base of the valley certainly retained year-round standing water because, even today, there is a perennial (if small) river in the valley bottom, and extensive remnants of river gravels below Sterkfontein and Swartkrans indicate that the river was larger and closer to the site in the past (Martini et al., 2003). There would have been a gallery forest which opened into woodland and possibly patches of grassland (Pickering et al., 2004).

## 3. Methods

In order to answer the question of how Member 2 formed, several methodological approaches were applied across different scales. At the macroscale, the chamber geomorphology and major depositional geometry were incorporated (e.g., Tankard and Schweitzer, 1976; Goldberg and Bar-Yosef, 1998; Latham, 1999; Farrand, 2001; Osborne, 2001; Sasowsky and Mylroie; 2004; Stratford et al., 2012). At the mesoscale, sediment facies descriptions and fabric observations in specific sections were correlated laterally and longitudinally through the Silberberg Grotto to associate specific features within the larger depositional framework (e.g., StW 573, solution cavities, colluvial, alluvial and collapse facies). At the microscale, thin section samples target key sections (see Fig. 1 for sample locations) and are described to provide sedimentological support for formation processes identified at the mesoscale. Sections chosen for meso- and microscale assessment (Figure 1) are associated with different areas of the Member 2 talus, which we consider to include all sediments that can be stratigraphically correlated with the StW 573 specimen, both upslope and downslope.

### 3.1. Cave Mapping

The existing plans (Martini et al., 2003) were unsuitable for the required sections so, a speleological topographic map was drawn of the main axes of the cave, which were then associated to the surface excavation site. Subsequently, a georeferenced, high definition 3D laser scan was conducted that integrates the main galleries of the Sterkfontein network. Special attention was given to the surface excavations (Stratford and Caruana, 2018), Silberberg Grotto, Elephant Chamber and Milner Hall areas, to ensure that the geometry of all possible parts of Member 2 were documented from their apex to the most distal parts. The profiles derived from these data are presented in the discussion.

### 3.2. Stratigraphy of the breccia

The processes of accumulation of Member 2 must be approached with a great caution. Depending on the origin of the sediments and their mode of accumulation, the associations with the fossils they contain can be misinterpreted. It is, therefore, important to know if Member 2 primarily accumulated through a gradual gravitational accretion of mostly surface-derived clastic material supplemented with sporadically collapsing blocks from the entrance or roof (Bruxelles et al., 2014; Stratford et al., 2017), or through a sudden introduction by a collapse or debris flows, for example with the collapse of an overlying gallery (Kramers and Dirks, 2017 a, b).

The specific stratigraphic context of StW 573 (Bruxelles et al., 2014) was studied prior to its dating by Granger et al. (2015). The objective of the previous study was to illustrate the diachronic relationships between the flowstone units associated with StW 573 and the Member 2 clastic deposits. Here this work is expanded to document the clastic units through the sequence of Member 2.

To understand the representative Member 2 sedimentation processes, four detailed stratigraphic sections were documented along the Silberberg Grotto Member 2 talus (as recognized by Partridge, (1978), Martini et al. (2003) and Clarke (2006) (Figure 1). The sections incorporate the eastern end of Member 2 and the proximal part of the talus on both flanks of its apex. Two further sections were documented in the central part of the western slope of the talus (proximal-medial). For each section, we describe each unit, its composition, organization, general fabric and stratigraphic association to the other recognized units, with particular attention paid to the organization of clasts and matrix (for mesoscale variables presented, see Table 1). Also noted are unconformities and stratigraphic irregularities (illustrated with dark bold lines between facies in Table 1).

### 3.3. Micromorphology

At the microscale, seven thin sections are presented that were strategically sampled through the exposed profiles of Member 2 in the Silberberg Grotto (see Figure 1) to better reveal the sedimentary structures that indicate the primary depositional processes of that unit. The placement of the samples was chosen to explore macroscopic stratigraphic correlations longitudinally and vertically through the proximal portions of both east and west flanks of Member 2 and the medial portions of the western flank Member 2, where facies could be stratigraphically associated with the StW 573. Three thin sections focus on facies within the proximal part of the eastern flank of Member 2. PE.A samples facies SA-5, PE.B samples SA-3, and PE.C samples SA-4 (Figures 2 and 3). One thin section, PW.A, focuses on a thick unit within the proximal west flank of Member 2, SB-5. Two thin sections focus on two facies identified to be stratigraphically associated with the StW 573 specimen - M.A samples SC-4 (B2b in Bruxelles et al., 2014), and M.B samples the underlying SC-3 (B2a in Bruxelles et al., 2014). One thin section, M.C, samples unit SD-3 of the SD profile and was taken slightly downslope of the SD profile for logistical reasons. The samples were embedded and thin sections prepared at the Thin Section Laboratory, School of Geosciences, University of the Witwatersrand (Johannesburg). They were then studied by Maire and Stratford in plane-polarized and polarized light. Microphotography was conducted at the Microscopy and Microanalysis Unit (MMU) at the University of the Witwatersrand using an Olympus BX63 Fluorescence microscope and automated sliding stage. Photographs were taken with a 4x objective under plane-polarized light (PPL) cross-polarized light (XPL). The primary objective of this approach was to provide microscale sedimentological support to the mesoscale observations and provide additional detail on the origin of the sediments. Detailed micromorphology and geochemical analyses are ongoing and will be presented elsewhere.

## 4. Results

### 4.1. Member 2 stratigraphy

The four detailed sections studied (Figure 1) allow the stratigraphy of Member 2 to be followed along the length of the chamber, incorporating StW 573 and highlighting the lateral and longitudinal facies variation along the slope. Facies characteristics from each section are summarized in Table 1 and described through their respective sequences below.

#### Section A - Silberberg Grotto, eastern end

This section (Figure 2) summarizes the observations made in the eastern end of the Silberberg Grotto where mining and blasting have exposed the complete depth of the unit across the width of the chamber.

From the bottom up, we have noted the following succession:

- **SA-1:** The base of the wooden staircase rests on a dark breccia comprised of blocks of chert and dolomite of all sizes within a sandy black-grey matrix cemented by calcite. This breccia corresponds to Member 1 (Partridge, 1978; Martini et al., 2003).

- **SA-2:** A calcite flowstone several decimeters thick conforms to the irregularities of the Member 1 breccia surface and embeds abundant blocks of chert. The abundance of blocks increases towards the top and becomes a chert breccia cemented by a calcite matrix. The top of this flowstone is relatively irregular and forms a dome in the central part of the section, descending along the irregular eroded flanks of Member 1. This represents the boss speleothem forming onto the clastic Member 1.

- **SA-3:** A reddish brown stony breccia overlies the flowstone in an angular unconformity. This level is not continuous in the section because it conforms closely to the topography of the underlying flowstone. The gradient is shallow, only a few degrees towards the East. At the eastern end, the unit divides into two. At the base, formed conformably on the SA-2 flowstone, a layer of about 10 centimeters consists mostly of bones embedded in an east sloping stratified, poorly indurated sandy-silty matrix with very low clast abundance. The abundant long bone fossils have been sorted but also, through deposition, are oriented in the direction of the slope (E-W).

Conformably overlaying this unit is about 10 centimeters of matrix-supported clasts and bones, which are organized in a planar fabric, although in some areas bones show linear fabric. This unit grades into a stony breccia with fragments of chert, dolomite, calcite concretions and bones.

- **SA-4:** The stony breccia is covered by a thick formation of coarse sandstones with infrequent dolomite and chert clasts. In detail, this formation represents decimeter-thick layers that can be followed along the length of the exposure. The granulometric sorting of these sandstones emphasizes the stratification of this formation and indicates some hydrodynamic sorting. Several darker layers that are very rich in microfaunal remains are also present (Figure 3). In the upper levels, discrete lines of clasts and several small layers of gravel can be seen.

- **SA-5:** Conformably overlying SA-4, coarse sandy facies continue. At about 40 cm thick, the unit includes very rare blocks of chert and dolomite. Here the bedding is less visible but those bones that can be seen have a planar fabric conforming to the dip of the bedded layers, demonstrating a succession of layers sloping eastwards.

- **SA-6:** Coarse sands grade into lighter and clearly laminated fine sands containing bones. Small layers of calcite are interbedded between some fine sand layers, indicative of an entrance in the process of closing.

- **SA-7:** The section is sealed by a finely laminated beige calcite flowstone. Continuous all along the exposure, this speleothem is visible throughout the upper eastern part of the Silberberg Grotto, where the above unit (Member 3) was deposited conformably onto it.

Such conformable deposition contrasts with the cavity-filling speleothem where formation of vertically downward forming speleothem fills a void (Bruxelles et al. 2014). This speleothem indicates a significant break in the clastic filling of the chamber and was identified by Partridge (1978, 2000) and Partridge and Watt (1991) as unit 3A. This speleothem provides a stratigraphic distinction between Member 2 breccia and the overlying Member 3.

#### Section B - Silberberg Grotto, west talus (proximal)

Fifteen meters down the western flank and topographically lower than Section A (Figure 1), the proximal section of the western slope of Member 2 has been exposed in a deep mining trench cut to access the thick speleothem covering Member 1.

From the bottom up, we have noted the following succession:

- **SB-1:** A partial concave casting of the thick Member 1-covering speleothem (the ‘boss’) is still preserved in the base of this unit’s breccia, which conforms to the steeply west dipping Member 1, not exposed anywhere west of this section.

- **SB-2:** To the east and at the base of the section, this breccia conformably rests against the boss. Where the breccia has formed directly onto Member 1, the contact is abrupt and erosional. Bedding is present and it is delineated by lines of chert gravel which are more abundant at the base. Dark gray in color, this unit represents the reworking of the altered dolomite and residual chert clasts of Member 1. SB-2 formed along a slope created by the initial conical accumulation of Member 1.

- **SB-3:** After an erosive unconformity developed following the general angle of repose of the SB-2, a coarse breccia with a reddish brown matrix-supported deposit formed. This 50 to 70 cm thick unit formed locally on the boss speleothem, whose imprint is still visible. The unit consists of an alternating sequence of coarse sands and small clasts within massive limo- sandy layers. Within sub-units, clasts retain a planar fabric. Some bones are visible in the exposure and are arranged according to the angle of repose of the slope.

- **SB-4:** Lying conformably above is a less indurated 30 to 40 cm thick generally relatively clast-poor unit supported by a sandy matrix. Bedding is noticeable in the upper part with discrete clast-rich lenses with consistent angles of repose. Differential induration of breccia within the unit also highlights the bedding.

- **SB-5:** The uppermost unit exposed in the section consists of about 60 cm of matrix-supported stony breccia that formed conformably onto SB-4. Discrete stony sub-units interstratify the silty-sand matrix. The base of this formation is clearly distinguished with a relative abundance of bones with strong fabric development according to the slope and orientation of the deposits. Remnants of SB-4 and SB-5 are preserved on the southern wall, opposite the Section B exposure and display the same sedimentological and fabric characteristics demonstrating the previous continuation of these units across the N-S width of the Silberberg Grotto.

- **SB-6:** The upper sedimentary reaches of the exposed section were mostly destroyed during mining. Between the current surface of SB-5 and the roof there could have been other levels of breccia. But these would not be very thick because a calcite flowstone which previously sealed this part of the deposit can be seen laterally, adhering to the roof only 20-30 cm above the surface of SB-5. From the remnant exposures, SB-6 is generally a fine reddish breccia of irregular thickness.

#### Section C - Silberberg Grotto, west talus (medial)

The section of the western talus published by Bruxelles et al. (2014) had focused on understanding the relationship between speleothem units and the StW 573-bearing breccia. Section C now focuses on the distinction of the different facies present in Member 2 in this area of the deposit, and it documents a new vertical exposure close to where StW 573 was discovered. Section C is advantageously orientated along an E-W axis of the gallery, longitudinally through the slope of Member 2 (consistent with the geometry of the clastic deposits) and in an area where the stratigraphy is less disturbed by the presence of filling speleothem units. To aid comparisons across space, the deposits distinguished are labelled with additional reference to those described in Bruxelles et al. (2014) (breccia (B) & calcite (F)). From the bottom up, we have noted the following succession:

- **SC-1:** At the current floor of the chamber, a stony breccia of pinkish color has been poorly exposed by mining activity and is difficult to describe in detail. It is a sandy matrix-supported breccia with poorly sorted clasts of a few centimeters to one decimeter in diameter.

- **SC-2:** The nature of the contact between SC-1 and the overlying finer, orange SC-2 unit is difficult to discern. Rich in fine clay/silt matrix, the unit contains some small scattered sub-rounded clasts and angular blocks. This breccia corresponds to B1 in Bruxelles et al. (2014).

- **SC-3:** Above, a relatively poorly cemented massive matrix-supported unit is present that is potentially secondarily altered. The poorly indurated nature of this unit has allowed the excavation of localized voids by water and the partial or complete secondary filling of those voids with calcite (B2a and F1, Bruxelles et al., 2014). The low, elongated shape of these voids, often sloping at thirty degrees towards the west, indicates the initial geometry of the unit. Clasts gradually become more abundant in the upper level of this unit and discernable decimetric blocks and smaller associated clasts conform to the same angle of repose of the voids and visible sedimentary contacts.

- **SC-4:** SC-3 grades into a clast-supported unit (B2b in Bruxelles et al., 2014) associated with the StW 573 skeleton and abundant elements of non-hominin primates and carnivores in fragmentary and complete condition. Poorly sorted and poorly organized medium to large (10cm) blocks are abundant and cemented within a reddish sand matrix.

- **SC-5:** The contact between SC-4 with SC-5 is locally disturbed by the presence of infiltrating calcite flowstones secondarily interstratifying the unit (F3 and F4 in Bruxelles et al., 2014). This brownish red silty-sand matrix-supported breccia with scattered small clasts and discernable bedding (B3 in Bruxelles et al., 2014) is of irregular thickness because it was partially destroyed by the mining of an overlying speleothem unit.

- **SC-6:** This speleothem unit seals the breccia deposits in this part of the Silberberg Grotto (F5 in Bruxelles et al., 2014). Despite being targeted by miners, it is still visible in section along the west gallery, where it fills the residual voids between the uppermost layers of breccia inclined to the west and the low roof. This unit represents a sealing calcite deposit which formed as a different unit to the cavity-filling flowstones, but it may be of similar geological age and formed under the same chamber conditions.

#### Section D - Silberberg Grotto, west talus (medial)

A final section was documented in the western end of the Silberberg Grotto (Figure 1). The section allows us to observe the continuation of recognized breccia units beyond and downslope of StW 573, confirming their continuity and the consistency of the sedimentary processes that governed their formation. Only the lower part of the sequence observed in Section C is exposed here because of the presence of a dolomitic pendant, the upper sediments of which were either eroded away or were removed through the mining process:

- **SD-1:** At the base of the exposed sequence (the floor of the chamber is not visible), a stony matrix-supported breccia is present with abundant medium to large (10 cm) blocks of chert and dolomite within a pink sandy matrix. Clasts display some fabric organization and indicate a dominant east to west slope. This unit corresponds to SC-1 of Section C.

- **SD-2:** An orange unit abruptly overlays SD-1 and is a finer, poorly consolidated matrix-supported deposit partially interstratified by infiltrating calcite speleothem. It corresponds to SC-2. As in SC-3, the shape and repose of the speleothem correspond to the shape of the voids they have filled, the latter being guided by the initial stratification of the unit and its differentially indurated internal bedding. Discrete lenses of small clasts are present and dip westwards, conforming to the overall geometry of the nearby units and stratigraphic contacts.

- **SD-3:** This matrix-supported bedded unit conformably overlies SD-2. At its base is a bed of poorly sorted sub-rounded to angular clasts that spreads from east to west with individual clasts showing strong fabric development, indicating an angle of repose of about 30 degree east to west.

- **SD-4:** This speleothemic unit fills the gap between the top of SD-3 and the bottom of a dolomitic pendant.

### 4.2. Thin sections

Although dedicated micromorphological analysis including geochemistry and mineralogy is ongoing, we describe here the major structures and features of the sediments and relate those to the prevailing formation processes involved in the development of that part of the talus and Member 2 in general. The location of each thin section sample in relation to the Silberberg Grotto and described sections can be seen in Figure 1. Below the general features are described before each sample description is presented from east to west across the Member 2 talus. We present annotated figures of PE.B (Figure 5) and PE.C (Figure 6) in plane-polarized and cross-polarized light to illustrate the general features representative of the samples. Figures of the other thin section samples are presented in the supplementary information.

**Figure 5:**
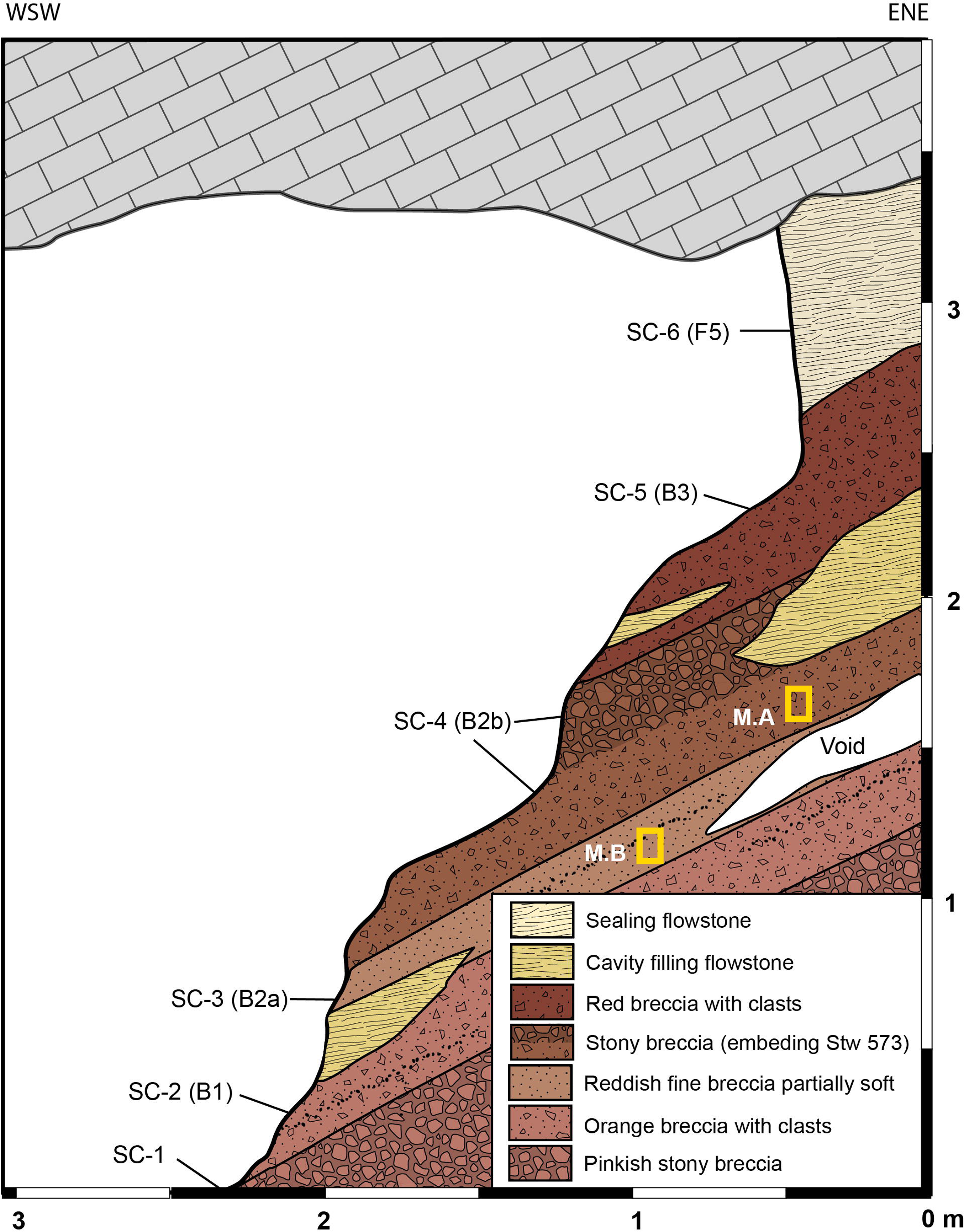
Synthetic representation of Section C (see Figure 1 for location).

**Figure 6:**
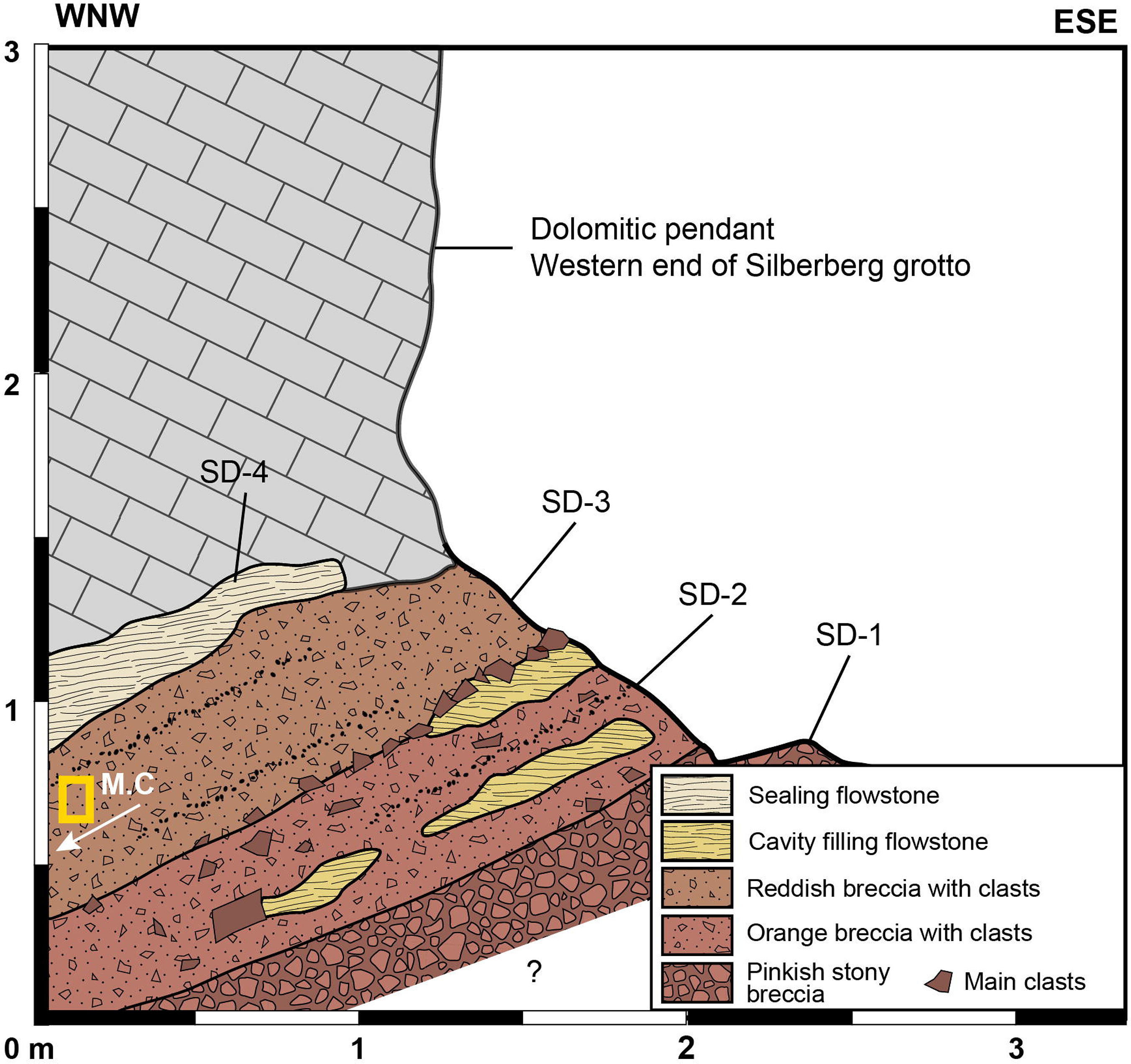
Synthetic representation of Section D (Western end of Silberberg Grotto, see Figure 1 for location).

#### General thin section features

##### Framework

Generally, the framework in all samples is an aggregated composition of infrequent small to medium clasts within an orange/red/brown and dark matrix of iron- and manganese-rich sandy silts that show varying levels of pedogenic aggregation. In situ decay/reworking has modified dolomites, broken down soil aggregates, and distributed manganiferous and ferruginous silts.

##### Structure

Structures are massive to stratified (e.g., PE.A, PE.C & PW.A) with occasional lamination and significant post-depositional calcite infiltration.

##### Clasts

These are: variably abundant, poorly sorted, angular to rounded chert clasts; small, sub-rounded to rounded dolomite clasts; small, rounded lateritic clasts with quartz grain inclusions; and rounded sandy pedogenic aggregates representing fragments of lateritic crusts. Clasts often have decayed surfaces and ferruginous coatings.

##### Matrix

The orange/red/dark brown matrix is generally massive and composed of coarse to fine quartz grains, and relatively abundant manganiferous and ferruginous lateritic silts deriving from pedogenic alterites that present as individual grains, aggregates and particle and pore space coatings. Variability in density is often due to variable induration of calcite.

##### Voids

Large void spaces are generally infrequent, elongated to rounded, dispersed and isotropic. Small voids are more abundant and form fine, irregular isotropic to linear networks occasionally directly associated with bedding, clasts and bioclasts. Voids are partially to completely filled with post-depositional calcite.

##### Postgenetic modification

Postgenetic void formation and filling with calcite has locally caused dispersion of sandy grains and iron and manganese-rich silts that fill voids and locally form coatings on clasts and individual particles. No distinct evidence is seen of in situ breakage or deformation through compression or collapse. Some post-depositional corrosion of chert and dolomite clasts is evident.

#### Individual thin sections

##### PE.A (Supplementary Figure A)

This sample is composed of: frequent poorly sorted angular, tabular and rounded chert clasts; sub-rounded to rounded dolomite clasts; rounded dark brown/black ferruginous and manganiferous clasts with quartz inclusions; and infrequent rounded sandy aggregates of pedological origin. Tabular and elongated particles tend to be horizontally or sub- horizontally orientated, perhaps indicating remnant structure and stratification. Most clasts have a dark Fe, Mn coating. The infrequent uncoated clasts represent contributions by autogenic cave breakdown. Some dolomite and chert clasts show heavy pedogenic weathering.

The orange/red matrix is generally massive and composed of fine grains of quartz, with small to medium-sized fragments of bone and silt, and is further indurated by the postgenetic calcite. Abundant manganiferous and ferruginous lateritic coating of grains occurs. Void spaces are frequent and larger voids are elongated and vertically orientated with partially or totally filled with calcite.

##### PE.B. (Figure 7)

This sample is composed of abundant void-filling calcite within a massive matrix, infrequent clasts that have little to no Fe, Mn coating, and abundant poorly sorted angular cancellous and cortical fossil fragments. Chert clasts are generally poorly sorted rounded to sub-rounded with notable presence of a large chert clast. Abundant rounded lateritic clasts (some with quartz inclusions) are present. Dolomite clasts are infrequent, rounded and decayed.

**Figure 7:**
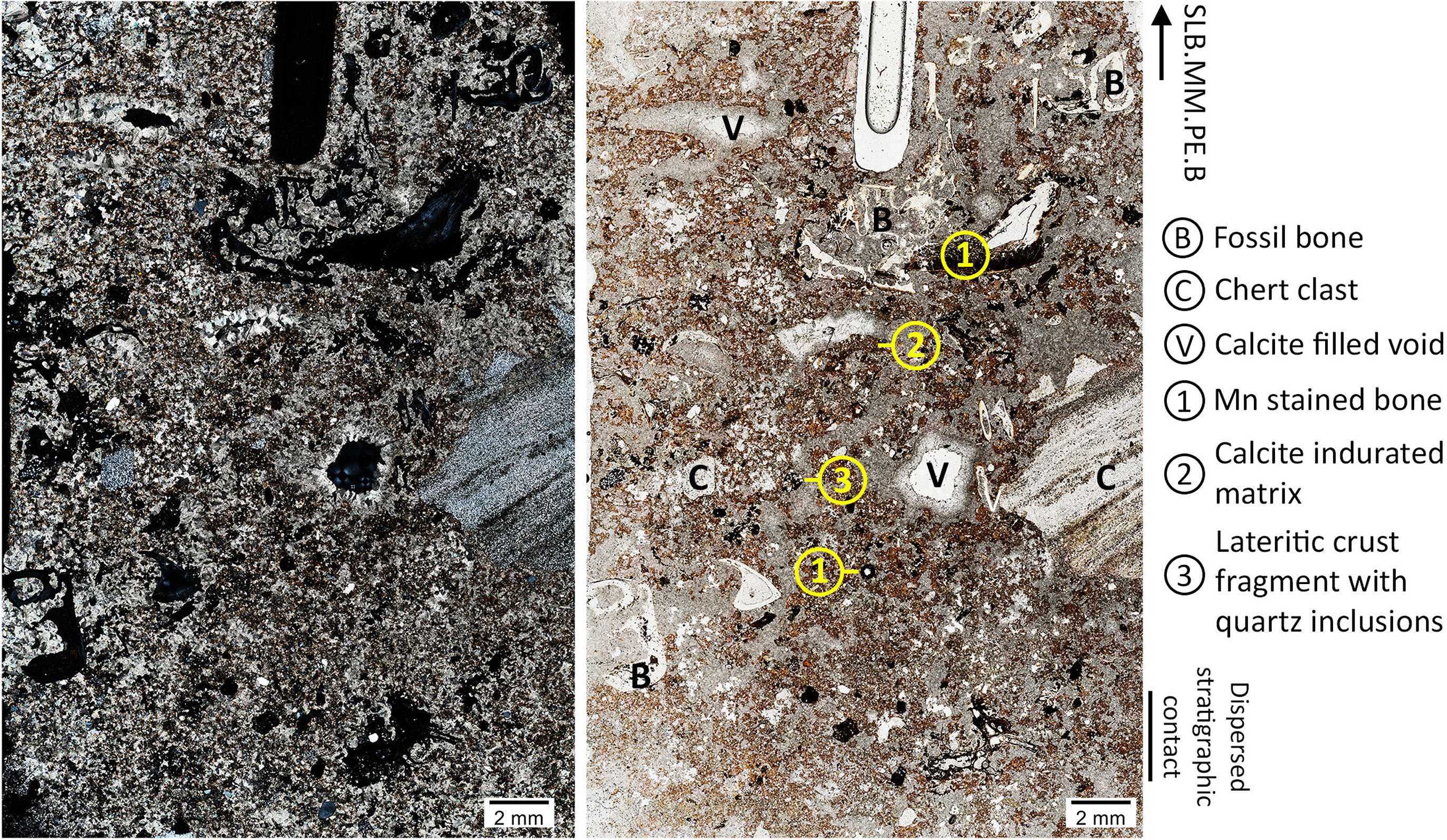
Sediment micromorphology sample PE.B (see Figure 1 for location) with key features and inclusions indicated. Right microphotograph is plane-polarised light, left microphotograph is cross-polarised light.

The orange/red matrix is generally massive and composed of fine grains of quartz and small to medium-sized fragments of bone within a clay and silt groundmass. Manganiferous and ferruginous silts fill voids and partially penetrate fossil fragments. Structure is unclear due to abundant postgenetic calcite.

Void spaces are abundant, varied in size and shape, and form irregular networks that pervade the sample and are frequently directly associated with clasts (including bioclasts) and decayed aggregated sediments. Voids are partially or totally filled with calcite.

##### PE.C. (Figure 8)

This sample is stratified sediment with few clasts. Small, sub-angular to sub-rounded clasts of dolomite (in various states of decay) and occasional large rounded bioclasts are distributed through the sample. Stratification is underlined by a large elongated void filled with sparitic calcite. Poorly sorted, rounded sediment aggregates are also present and more abundant in the lower half of the sample. Below this level, small, sorted angular sand-sized quartz grains are associated with generally horizontal strata and have been dispersed by postgenetic void formation and calcite filling.

**Figure 8:**
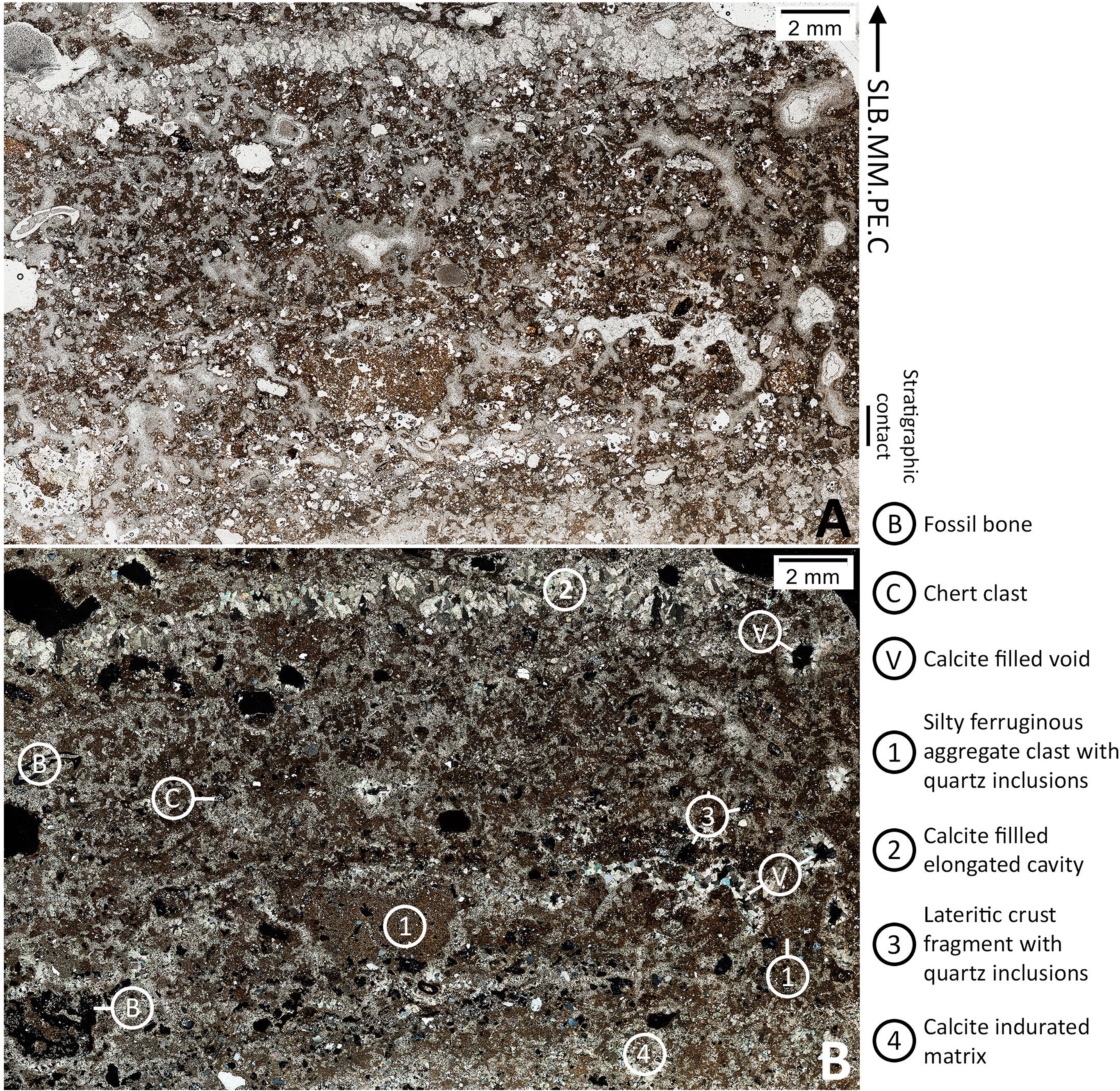
Sediment micromorphology sample PE.C (see Figure 1 for location) with key features and inclusions indicated. Upper microphotograph is plane-polarised light, lower photograph is cross-polarised light.

The orange/red matrix is generally massive between strata and composed of fine grains of quartz and small to medium-sized fragments of bone within a silty clay ferruginous groundmass. Manganiferous and ferruginous silts are more abundant in the upper half of the strata as lateritic grains, silty grain coatings, fill voids or adhere to poorly preserved sediment aggregates. Matrix is horizontally bedded.

Voids are infrequent, occasionally rounded, but generally form irregular networks that spread horizontally across the sample conforming to the bedding of the sediment.

##### PW.A. (Supplementary Figure B)

This sample is composed of: infrequent, poorly sorted sub-angular to sub-rounded chert clasts; small, sub-rounded to rounded dolomite clasts; and infrequent rounded lateritic aggregates of pedogenic origin with small quartz grain inclusions. Chert clasts are partially coated with manganiferous and ferruginous silts and generally restricted to the upper surface. Clasts are better sorted and more abundant in the upper reaches of the sample.

The orange/red/dark brown matrix is generally massive and composed of coarse to fine quartz grains, and relatively abundant manganiferous and ferruginous lateritic silts present as individual grains, aggregates and particle and pore space coatings.

Large void spaces are infrequent, round in shape and irregularly dispersed through the sample. Small voids are abundant and characterized by fine irregular networks extending roughly horizontally across the sample. Voids are both unfilled and filled by calcite and some show possible generational dissolution and deposition of calcite forms.

##### M.A. (Supplementary Figure C)

The framework is an aggregate composed of clasts of poorly sorted angular (few) to smaller rounded chert clasts (2-5mm) and ferruginous, rounded indurated red-orange grains (2-10mm) of sandy sediments of pedogenic origin and irregular dark brown to black ferruginous and manganiferous lateritic grains with few quartz inclusions. Dolomite clasts are rare, small and rounded. Clasts show no organization and are not directly associated. Iron and manganese coated laterite fragments are also observed associated with angular to rounded poorly sorted mono- and poly-crystalline quartz grains.

The orange/red ferruginous matrix is composed of fine grains of quartz and fragments of bone. Silty sand matrix is further indurated by the postgenetic calcite, with partially or totally calcified voids. Void spaces are generally small and widely distributed and larger voids are elongated and isotropic.

##### M.B. (Supplementary Figure D)

This is a generally massive sediment composed of fine sandy and millimetric fragments of chert with ferruginous soil cutans. Clasts are small, scarce, poorly sorted, rounded to sub-rounded chert and sandstone with Fe and Mn coatings. Infrequent small rounded dolomites are present in various stages of decay. Clasts show no organization and are not directly associated. Structurally, the sediments are less aggregated than in M.A. Some grains show very dark coatings of manganese and ferruginous grains of orange-red goethite type are present.

The orange/brown matrix is composed of fine quartz grains and ferruginous and manganiferous silts and clays penetrated by a cementation of white calcite. Large void spaces are generally infrequent, elongated to rounded, and restricted to the upper portion of the sample, dispersed and isotropic. Small voids are abundant and are not filled with calcite.

##### M.C. (Supplementary Figure E)

Two layers are present: a dark, upper layer and a lighter lower. The upper layer is composed of poorly sorted angular to rounded chert clasts, the majority of which has Fe and Mn coatings (some more angular chert clasts have no coatings at all), occasional well-preserved bone fragments and infrequent grains composed of ferruginous lateritic silts and clays with rounded poorly sorted quartz inclusions. Clasts are more abundant in the upper part of the upper unit.

The upper brown-red unit matrix is generally massive, heavily indurated and composed of fine quartz and iron and manganese-rich groundmass with no clear aggregates present. Voids are abundant and calcite fills larger elongated and perpendicular void networks with a strong left to right inclination. Small voids pervade the sample and almost all voids are filled in the upper unit.

The lower unit is clast-poor and heavily indurated and shows a “chopsticks” of manganese crust. Clasts are sorted, rounded to sub-rounded and made up of chert and quartz. Some remnant sediment aggregates may be present. The unit is dominated by white sparitic calcite with a low density of clay-ferruginous matrix and small crust Fe/Mn grains. Many small, unfilled voids are visible throughout the matrix with a possible role of a dissolution phase.

## 5. Interpretation

### 5.1. Global geometry of Member 2 deposit from macro- and mesoscale data

The geometry of Member 2 in the Silberberg Grotto indicates the deposit forms a single cone talus, the apex of which is located in the eastern end of the chamber (Figure 1) beneath an original entrance which fed the chamber and is now choked. On the eastern flank, the east-dipping Member 2 is sealed by a calcite flowstone (SA-7 here; Unit 3A in Partridge, 1978) separating it from the overlying breccia (Member 3 in Partridge, 1978). The western flank of the Member 2 talus is clearly observable since its exposure from extensive speleothem mining in the central and western areas of the chamber. The western flank slopes at a 30 to 40 degree gradient westwards towards the Elephant Chamber for almost thirty meters before exiting the Silberberg Grotto through three passages.

The general shape of the talus is partly influenced by the underlying Member 1 breccia and the overlying boss speleothem, which here we consider to be part of the autogenic Member 1 formation process. During the vadose collapse of the cave, a process that formed Member 1, the thickest Member 1 deposits formed where there was greatest decay of the dolomite, i.e., where faults in the dolomite are most abundant on the southern boundary of the system (Stratford, 2017). The highest part of the roof of Silberberg Grotto is in the east, the area where the largest amount of dolomite and chert material was removed to the base of the chamber during the vadose, joint-governed collapse, thereby forming a talus of chert blocks encased in a black-gray sandy matrix subsequently cemented by calcite (i.e., Member 1 and the associated boss speleothem). The collapse of the roof in the eastern area of the chamber also thinned the overlying host rock (relatively), enabling increased vertical water movement, precipitation of speleothem and ultimately the potential for the development of an entrance above the Member 1 talus. All erosive and depositional processes found in the Silberberg Grotto originate from this location. The eroded western flank of Member 1 dips more steeply than the eastern flank, a morphology that has partially controlled the distribution and resulting asymmetrical geometry of Member 2 with sediments preferentially being deposited to the west (Sections B, C and D). The eastern flank (Section A), shallower in repose, received less sediment through fluvial processes as observed by Partridge (1978) and Clarke (2006) and evidenced by the sorted and well-orientated fossils exposed in the eastern breccia wall.

As observed by previous researchers (e.g., Partridge, 1978; Martini et al., 2003), the Member 2 talus geometry indicates that the deposit’s entrance is located vertically to the apex. At the macroscale, surveys from the Silberberg Grotto to the landscape surface (Figure 9) revealed that there is no dolomite above the highest point of the talus and the remnants of a vertical conduit is present, still filled with breccia, and located in the Member 4 south area of the surface excavations. The upper contact of Member 3 is an important feature, which, and at this juncture, we don’t have the exposures to document. From inside the Silberberg Grotto, Member 3 ascends nearly vertically to the ceiling, its morphology governed by the shape of the boss speleothem it formed around. The lower contact of Member 3 is clearly distinguished from Member 2 by an intercalating flowstone (SA-7 in Figure 3; Unit 3A in Partridge (1978) and Partridge and Watt (1991)). On the western talus a speleothem unit sealed Member 2. Much of this was mined away in the proximal sections of the talus as it formed close to and in places onto the boss speleothem but we recognise it as correlating with SA-7, SC-6 and SD-4. In section B, remnants of this may adhere to the ceiling above the mined unit (with remnants preserved) SB-6. This speleothem has been sampled for palaeomagnetic dating and was found to yield a normal magnetic polarity (Partridge, 1999; Herries and Shaw, 2011). If the cosmogenic dates of Granger et al. (2015) are correct then this speleothem formed under normal global magnetic polarity may relate to the Gauss-Gilbert boundary, dated to 3.58 ma (REF).

**Figure 9:**
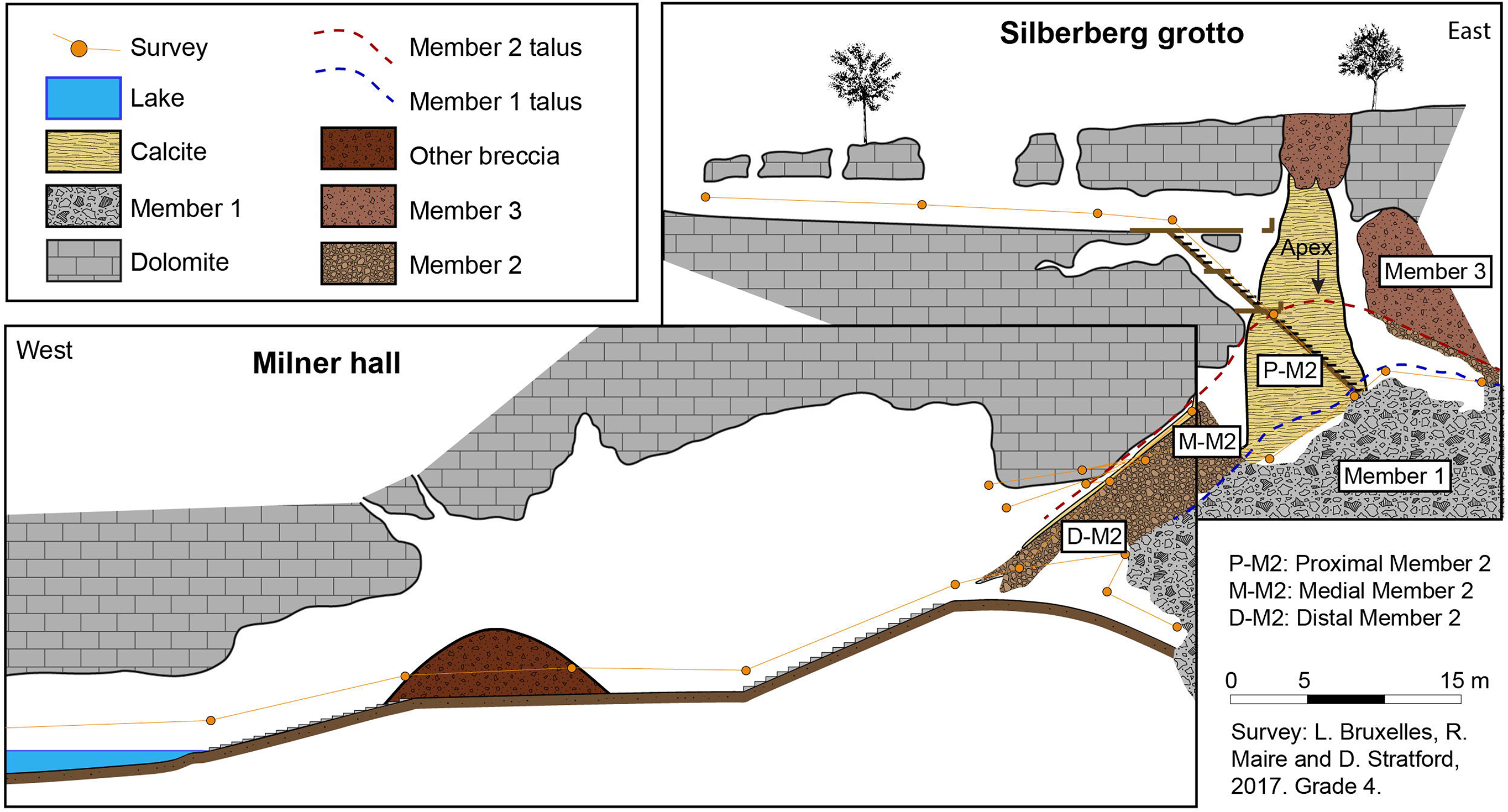
Global section along the Member 2 talus, from Silberberg Grotto to the Milner Hall.

The original entrance to the chamber which fed Members 2 and 3 looks to have been choked by Member 3 (Figure 9). The highest point of the roof here is breccia. Immediately above this point, on the surface, is the southern-most extent of a solution cavity-riddled breccia that is called Member 4 (Figure 1). We therefore do not observe a contact between Member 3 and Member 4 in this area. It may be that Member 3 is very thin here, the contact is near vertical (as is often the case close to walls and entrances), or the contact has been mixed. The nature of the upper contact of Member 3 is therefore crucial to our understanding of the nature of Member 3.

The general shape of the talus, the inclination of the slope (between 30 and 40 °), and the longitudinal profile of the slope, which is slightly concave towards the base, are indicative of a colluvial talus (Kirkby & Statham, 1975; Statham, 1976; Postma, 1986; Brochu, 1987; Perez, 1989; Saeter, 1998; Bertran and Texier, 1999; Bertran et al., 1995 and 1997). A proximal part, located at the apex, a medial part which is visible all along the western part of the Silberberg Grotto, and a distal part can be observed. The distal part, more difficult to observe because of mining damage, can be tracked into the Elephant Chamber and Milner Hall. The macro- and mesoscale mapping demonstrates that the slope of Member 2 is continuous and coherent from apex to the Elephant Chamber and the Milner Hall, the implication being that the whole depth of the system was formed prior to significant opening to the surface, and early Member 2 sediments accumulated toward the base of the system (as suggested by Wilkinson, 1973, 1983; Stratford, 2014). This challenges the karstification model suggested by Partridge (1978), Partridge and Watt (1991), and Pickering and Kramers (2010).

In the eastern part of the Milner Hall, a stony breccia remains cemented against the south wall with a preserved surface inclined at 38 ° westward. This heavily undercut deposit can be followed for about 20 meters, from the eastern end of Milner Hall East up a continuous slope through a narrow passage articulating with the Silberberg Grotto about 10 m below where StW 573 was discovered. It indicates that the breccia in the Milner Hall has the same geometry as Member 2 and represents a lateral and longitudinal continuity of Member 2 (Figure 9).

### 5.2. Member 2 breccia accumulation

The mesoscale and microscale evidence collected through Member 2 makes it possible to identify varying modes of talus accretion. This is a crucial step to understanding the taphonomy of the faunal assemblage, but also the association between sedimentary features and interred fossils, particularly StW 573, whose association with the nearby flowstones and sediments has been the subject of debate concerning the dating of the specimen (e.g. Clarke, 2002b; Pickering & Kramers, 2010; Granger et al., 2015; Kramers & Dirks, 2017a,b; Stratford et al., 2017).

First, it is clear that Member 2 is stratified through its entire thickness along the length of the talus (sections A to D, Figure 1). Whether east or west of the entrance, several successive breccia units can be distinguished and linked to the structures and features found in the thin sections which present in all samples variable abundances of the same composition of primarily allochthonous materials with varying degrees of modification and infrequent inclusion of autochthonous clasts. All thin sections indicate the gradual erosion of well-developed lateritic soils on the surface (as suggested by Stratford et al., 2014 from sediments attributed to an early distal Member 2), contributing an iron-rich sandy matrix composed of a variety of soil aggregates, well weathered chert clasts and fossil fragments. Autochthonous clasts are un-coated, fresh and angular and do not represent the dominant clast contribution in any sample.

In Section A, the earliest infilling of breccias are documented, covering the Member 1 breccia and the boss speleothem. The stony and bone-rich breccia (SA-3) molds the irregularities of the Member 1 dome. It constitutes the very first level of allochthonous breccia coming from the outside and is the result of the opening of an entrance above the apex of the talus, where the vault was the highest. This level of stony breccia (SA-3) could correspond to SB-3 in the proximal medium talus and the unit partly mined out at the base of Section C (SC-1 and SD-1) (Figure 10) - with finer composition generally associated with more proximal facies of the same unit.

**Figure 10:**
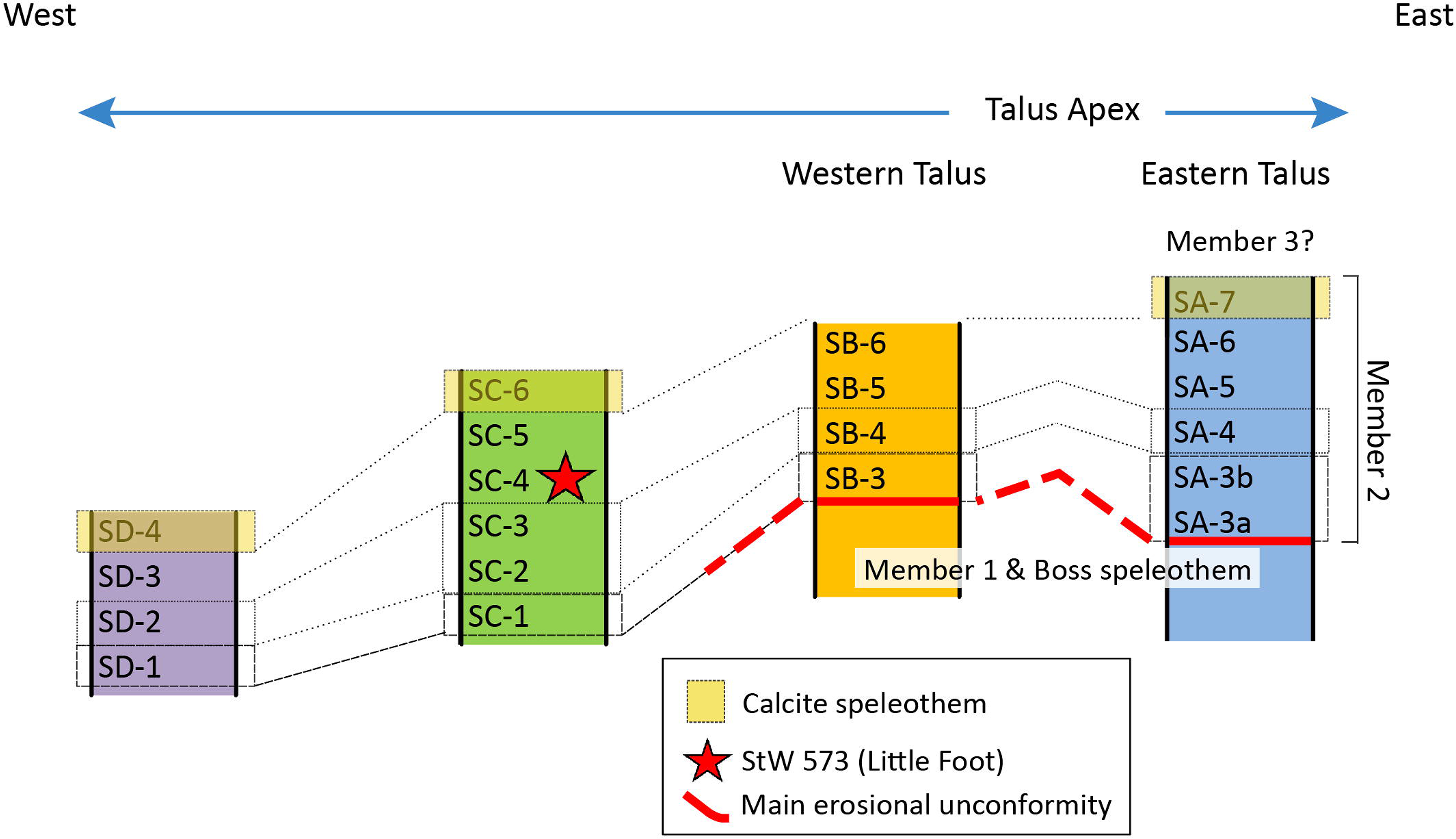
Proposition of correlations of the different sedimentary layers through the Member 2 talus in the Silberberg Grotto.

Above, still in section A, the unit SA-4 is coarse breccia of variable thickness. This breccia is also stratified and shows a progressive accumulation by fluid run-off, which alternates between detrital layers coming from the erosion of the surface soil and beds rich in microfauna. SA-4 can be correlated to SB-4 and SC-3 (and perhaps SC-2) further downslope to SD-2 (Figure 10), which demonstrates an increase in clast size and abundance due to longitudinal sorting. A tentative correlation can also be made between SB-5 and SC-5, suggesting a discrete development of SC-4, or perhaps to SC-4 & SC-5, suggesting a thickening of SB-5 downslope. Across these sections we find the same facies successions, all in geometric coherence with the layers that can be connected following a slope of 30 to 40°.

Along the slope, the particle size of the clasts and the proportion of matrix vary from the proximal portion to the distal portion of the slope. This is entirely consistent with the formation processes found on a colluvial accumulation in a talus (Statham, 1976; Kirkby & Statham, 1975; Brochu, 1987; 1986; Perez, 1989; Saeter, 1998; Bertran and Texier, 1999; Bertran et al., 1995 and 1997; Saeter, 1998; Bruxelles et al., 2017). Proximal facies are characterized by finer sediments and generally lower abundance and smaller sizes of clasts. The distal facies of the slope are characterized by large blocks, most often without fine matrix, and the voids between the clasts can be subsequently filled by reworked sediments and speleothem. In the medial part of the talus, facies distributions are variable and discrete stony layers can be locally preserved - like SC-4, the unit in which Little Foot was found. The longitudinal stratigraphic correlations along slope and the associated variability in clast abundance and size indicate that Member 2 was deposited primarily through gradual autochthonous colluvial processes from a slowly eroding landscape covered in potentially well-developed lateritic soils (Moeyersons and De Ploey, 1976; Osborne, 1978 and 2001; Mihevc et al., 1998; Latham, 1999; Pederson et al., 2000; Klimchouk, 2006; Martini et al., 2003; Martini 2001; White, 2007; Bruxelles et al., 2017) through the high aven-like entrance above the eastern end of the chamber. Deposits were added to sporadically by relatively slow breakdown of the walls and the vault.

### 5.3. Taphonomy of StW 573

A detailed account of the taphonomy of the StW 573 specimen is provided by Clarke (2018, this issue) and follows previous observations presented in Clarke (2002b). In addition to this, here the position of the StW 573 body is discussed in relation to the sedimentary history of Member 2. Importantly, StW 573’s body is orientated parallel to and at the angle of repose of the underlying and overlying deposits. It can be proposed that body fell onto the apex of the slope, directly beneath the opening in the eastern end of the chamber, and rolled perpendicular to the slope for 10 m down the western flank. Rotation of the body to a perpendicular orientation stabilized the body on the slope and the specimen’s leg passed over the right as it came to a stable position with the left arm upslope above the body. Post-depositional modifications are all local events with no evidence of mass movements of stone or matrix observable in the specimen’s preservation or associated sedimentary units above, below or laterally. Localized breakage can be limited to single events (although not necessarily at the same time) of isolated impact, compaction or subsidence, with broken bones, culprit clasts and voids, and associated fragments all remaining associated until excavation. The body position, mummification (Clarke, 2002b) and localized vertical disruption of only the torso of the specimen (into the cavity below, as opposed to downslope movement) all indicate a solitary deposition of the specimen, a gradual burial and isolated post-depositional modification, all correlating with the general sedimentary history of Member 2.

## 6. Discussion

New multiscale research presented above provides the sedimentological and geomorphological evidence to scrutinize previous Member 2 breccia formation hypotheses more closely. Below, the hypotheses are discussed in relation to the evidence presented above.

An age of 2.8 - 2.2 Ma for StW 573 and Member 2 was proposed recently by Pickering and Kramers (2010) and Herries and Shaw (2011). The dating of flowstones below the Silberberg Grotto by Pickering and Kramers (2010) which yielded relatively young ages supports the Member 2 epiphreatic karstification formation model proposed by Partridge (1978) and Partridge and Watt (1991). In this model, Member 2 represents the basal allogenic sedimentary deposit at Sterkfontein, with a series of deeper passages and chambers forming later and being filled by younger or reworked sediments. Macroscale mapping of Member 2 has provided crucial clues to test this hypothesis. This work instead confirms suggestions by Wilkinson (1973, 1983) and Stratford et al. (2014) that Member 2 previously extended beyond the Silberberg Grotto into the deeper caverns of the Elephant Chamber and the Milner Hall. Distal remnants matching the geometry of Member 2 can be found adhering to the roof and walls of the eastern Milner Hall and can be traced close to the current base level. The sediments represented there are characteristic of distal facies of colluvially accumulated deposits with 30-40° gradients, consistently demonstrated in the bedding of Member 2 both inside and outside the Silberberg Grotto. This evidence increases the vertical and longitudinal distribution of Member 2. If basal distal sediments are present on the floor of the Milner Hall as suggested by Stratford et al. (2014), another four meters vertical depth can be added - a cumulative depth of 19 m for Member 2 sediments).

Micromorphological evidence suggests the slow accumulation of well-developed lateritic soil aggregates and relatively low energy post-depositional modification of those soil structures. Structurally, the facies are similar to colluvial cones associated with slowly accreting ‘overland flows’ (Bertran and Texier, 1999), although subaerial modifications (e.g., rain splash) are limited in this context and post-genetic modification and void development differ due to the calcite intrusion. A limited catchment area of sediments around a relatively small opening near the top of the Sterkfontein hill also suggests a relatively limited source of sediments and low sedimentation rate. Mesoscale documentation of facies through the Silberberg Grotto allows us to correlate the sedimentary units longitudinally. Associating the identified facies variation longitudinally and vertically through the deposits allows the unification of the entirety of Member 2 into a single sedimentary and spatial framework. It is clear that colluvial processes dominate facies development and variability, forming coarser facies at the distal end and finer facies at the proximal end (Figure 10).

Considering the increased lateral and vertical distribution of Member 2, the low frequency of autogenic clast contribution, and the limited sediment source (topographically), we suggest this evidence supports Granger et al.’s (2015) hypothesis of a slowly denuding landscape and slow accretion of sediments as a gravitational colluvial talus, which developed into a spatially extensive Member 2 deposit, within which StW 573 was buried close to the end of the unit’s accumulation. In response to Granger et al. (2015), Kramers and Dirks (2017a, b) suggested that the stratigraphic and chronological association between StW 573 and Member 2 may be more complex. They posited that StW 573 and the sediments associated with it were subjected to a ‘two-stage burial scenario’, with StW 573 collapsing from a chamber above that is now eroded away. The mixing of sediments and the Little Foot specimen would constitute the formation of a ‘secondary’ deposit and therefore should be evident in the structure and composition of the deposits around StW 573. While we recognise that the original entrance to the Silberberg Grotto may now be where M4 south is, the entrance having been choked by Member 3, the location of the collapse proposed by Dirks and Kramers (2017a) is beyond the known eastern extent of the Silberberg Grotto and is east of the Member 2 talus apex (Stratford et al., 2017). Localised collapse above the western end of the chamber would be the only process that would enable the secondary deposition of a completely articulated skeleton. In addition, collapse deposits in caves are generally characterized by increased abundance of angular autogenic clasts, isotropic fabrics and fragmented speleothems and fossils. Specific reply was given by Stratford et al. (2017) regarding the macroscale and taphonomic implications of such a scenario, but further investigation was needed to clarify the association of the sediments close to StW 573 with the broader depositional framework to test their hypothesis, which we have done here.

Structurally, no evidence was found suggesting localized collapse in the vicinity of the specimen. SC-4 (B2b in Bruxelles et al., 2014) conforms to the geometry and organization of the bounding units and can (although tentatively at this point) be correlated to the more proximal unit SB-5. Voids, clasts and bioclasts below, within and above conform to a consistent 30 - 40 ° east to west slope suggesting regular bedding of all the Member 2 units including SC-4. In thin section, samples M.A and M.B show no composition difference to the other samples in terms of abundance of autogenic clasts and no distinct structures indicative of collapse microfacies. The generally low abundance of angular autogenic particles suggests slow internal chamber and entrance breakdown. The absence of fragmented speleothem, a common indicator of collapsed chambers, is absent at the meso- and micro-scales. The sediments have developed through the same processes as the other sampled units throughout the deposit. Taphonomically, although StW 573 is the only articulated full skeleton in Member 2, articulated and antimeric elements are found in the abundant fauna closely associated with the specimen, and as discussed above and by Clarke in this issue, post-depositional modification of the specimen is limited to localized and isolated low energy events affecting individual parts of the articulated skeleton, with each modification being preserved in its primary modifying context (i.e., bones, clasts, voids, and fragments are still associated). The articulation of the left lower arm and hand is an excellent example of this, with clasts still associated at the breakage points of the radius and ulna and slightly displaced wrist bones. This would not be expected in a collapse scenario with turbulent movement of clasts, bones and sediments. It would also not be expected if the mode of initial burial or subsequent movement were through debris flow mechanisms, where a mass movement of clasts and sediments create turbulent flows and clast-supported lateral and longitudinal lobes (e.g., Van Steijn and Coutard, 1989; Bertran and Texier, 1994; Major, 1998; Jameson, 1999; Bertran and Coussot, 2004; Jakob, 2005). Sedimentary structures indicative of debris flow processes are not evident in any of the Silberberg Grotto Member 2 exposures.

## 7. Conclusions

The multiscale evidence presented above has sought to test recent hypotheses regarding the stratigraphic and chronological history of Member 2 and the StW 573 specimen. Utilizing these tools provided the opportunity to clarify not only the history of the infilling of the Silberberg Grotto, but also the source and processes of accumulation of the Member 2 sediments and their association with the StW 573 skeleton. We do not provide new dates for the specimen but attempted to find supporting stratigraphic and sedimentological evidence to test the hypotheses put forward by Pickering and Kramers (2010), that some of the sediments in the western Silberberg Grotto may not be Member 2 and that the deposit is restricted to the Silberberg Grotto (in support of proposals by Partridge (1978) and Partridge and Watt (1991) for an epiphreatic karstification process forming the caves).

Our work has also addressed Kramers and Dirks (2017a,b) recent proposal that a single sample included in the nine sample cosmogenic nuclide isochron published by Granger et al. (2015) could indicate reworking of sediments and a two-stage burial scenario for the Little Foot skeleton involving a collapse from an upper chamber. We have drawn upon the most recent stratigraphic work conducted by Bruxelles et al. (2014) and absolute dates for Member 2 and the StW 573 specimen proposed by Granger et al. (2015).

The sequence of deposits associated with ‘Member 2’ (as originally identified by Partridge, 1978) can coherently be linked to the evolution of the Silberberg Grotto and includes the formation of the chamber, the opening of the entrance high in the roof, geometric constraints of the first allogenic infillings, episodic sedimentary hiatuses, differential sedimentary processes conforming to the geometry of the chamber, and final infilling and precipitation of the sealing calcite speleothem which represents the end of the Member 2 sequence. Throughout the sequence a consistent sedimentary framework is identified, which typifies many of Sterkfontein’s deposits, characterized by slow accretion through colluvial talus development. The consistent bedding, concave longitudinal shape of Member 2 and longitudinal facies development are all characteristic of this mode of accumulation and attest to a slow accumulation with no distinct collapses. Stratigraphic and facies correlations can be made that unify the deposit from its eastern flank exposures through to the remnants adhering to the walls and roof of the Milner Hall, confirming its previously proposed extent close to the current base level (Wilkinson, 1973, 1983; Stratford et al., 2014). The extension of Member 2 into the Milner Hall thus challenges the epiphreatic karstification model proposed by Partridge (1978) and Partridge and Watt (1991), suggesting the complete depth of the caves as we see it now had been developed at the time of its opening to the surface and that the deepest chambers may contain the oldest deposits.

Microscopically, the similarities between samples indicate the same general accumulation process and suggest a slowly eroding landscape with allogenic sediments contributed from well-developed lateritic soils and slow autogenic cave breakdown but extensive localized calcite modification. A distinct lack of dolomite clasts and low frequency of uncoated, angular chert clasts support a gradual mode of accumulation with no obvious collapse facies found through the sequence. The relatively gentle stratigraphic history of Member 2 supports the taphonomic history of not only the StW 573 specimen, but also the abundant other faunal remains which remain articulated and free from extensive biogenic or post-depositional modification. In conclusion, the evidence presented here supports the long development of a previously very large Member 2 and challenges the formation hypotheses made by Pickering & Kramers (2010) and Kramers & Dirks (2017a) and the associated implications for the alternative younger age for StW 573.

## Supporting information

## Acknowledgements

The support of the AESOP+ program, the Claude Leon Foundation, the DST-NRF Center of Excellence in Palaeosciences (CoE-Pal), The Palaeontological Scientific Trust (PAST), the French National Institute for Preventive Archaeology Research (Inrap), the Embassy of France in South Africa and the French Institute of South Africa (IFAS) towards this research is hereby acknowledged. Major funding for the Sterkfontein excavations has been provided by National Research Foundation grants to KK (#82591 and 82611) and to DS (#98808) and by PAST, without whose support this research could not have been able to continue. We thank Andrea Leenen and Robert Blumenschine for their help in securing major corporate funding, including sustained support from Standard Bank and JP Morgan. Opinions expressed and conclusions arrived at are those of the authors and are not necessarily to be attributed to the Center of Excellence in Palaeosciences.

---

Table 1: Summary of structural and sedimentological characteristics of the identified stratigraphic units within each documented section. See Figure 1 for location of documented sections.

## Supplementary materials caption

Supplementary Figure A. Sample PE.A (see Figure 1 for location) microphotographs in plain- polarised (left) and cross-polarised light (right). Arrow indicates orientation of sample and scale is shown on each image.

Supplementary Figure B. Sample PW.A (see Figure 1 for location) microphotographs in plain- polarised (left) and cross-polarised light (right). Arrow indicates orientation of sample and scale is shown on each image.

Supplementary Figure C. Sample M.A (see Figure 1 for location) microphotographs in plain- polarised (left) and cross-polarised light (right). Arrow indicates orientation of sample and scale is shown on each image.

Supplementary Figure D Sample M.B (see Figure 1 for location) microphotographs in plain- polarised (left) and cross-polarised light (right). Arrow indicates orientation of sample and scale is shown on each image.

Supplementary Figure E. Sample M.C (see Figure 1 for location) microphotographs in plain- polarised (left) and cross-polarised light (right). Arrow indicates orientation of sample and scale is shown on each image.

## Footnotes

The term ‘breccia’ is often misused to refer to any and all cemented or decalcified cave infill in the Cradle of Humankind sites. The sediments classified as ‘breccia’ are highly variable and do not conform to the formal sedimentological definition of a breccia. Due to the pervasive use of the term in the COH literature, we have continued with its traditional use to describe any cave infill calcified or decalcified.

