## Supplementary figures and images for "A multiscale stratigraphic investigation of the context of StW 573 ‘Little Foot’ and Member 2, Sterkfontein Caves, South Africa"

### Supplementary file 1

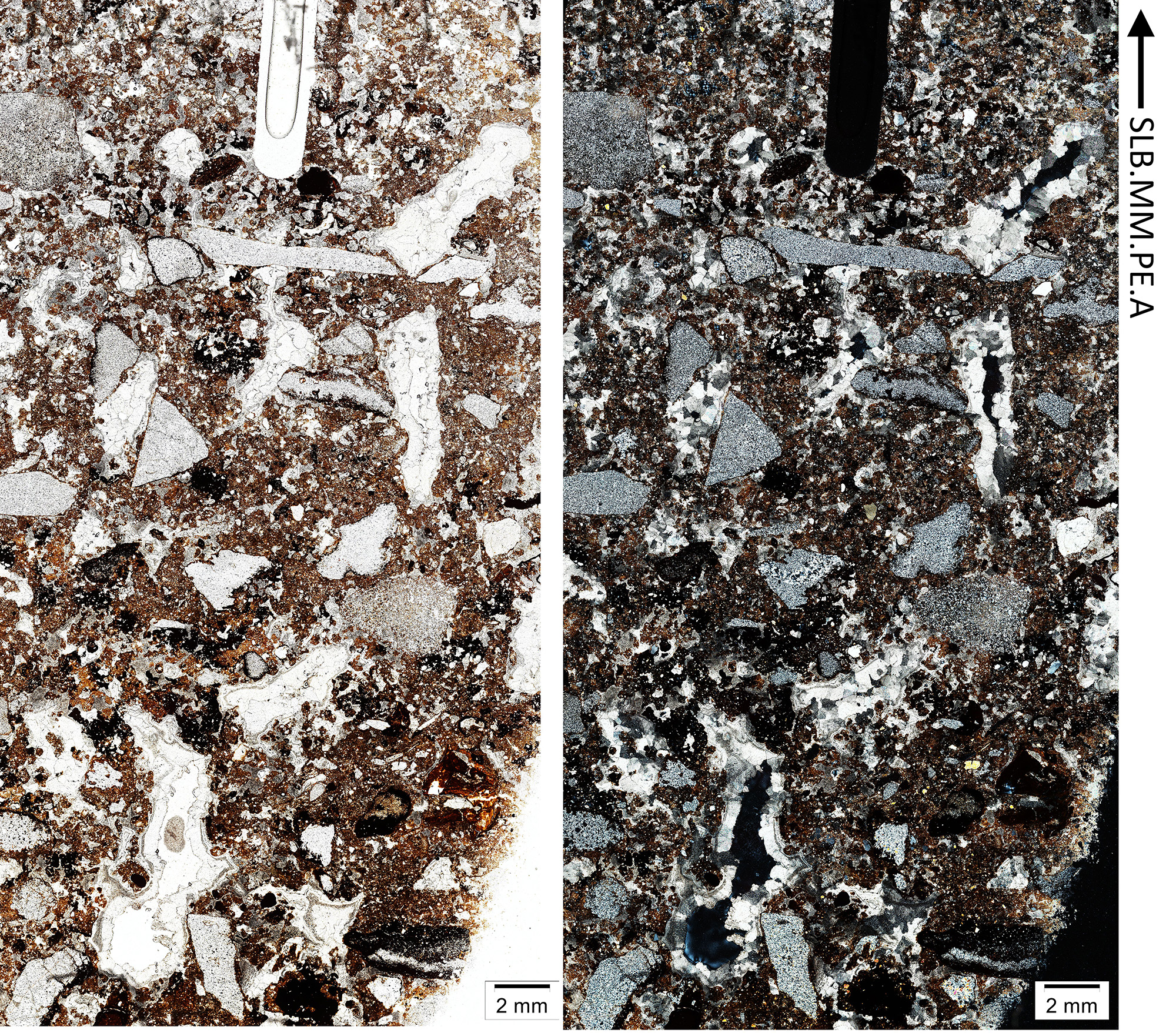

### Supplementary file 2

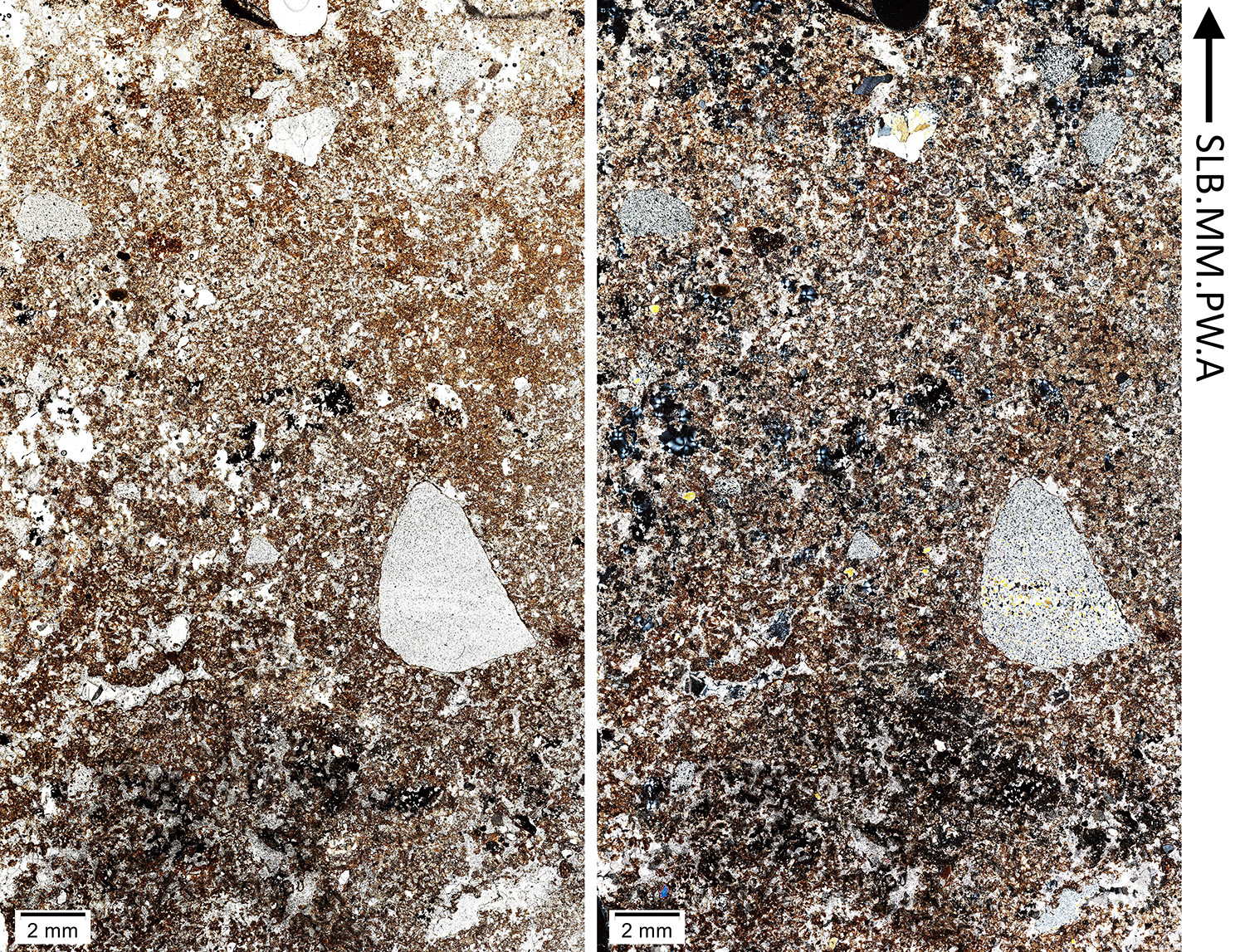

### Supplementary file 3

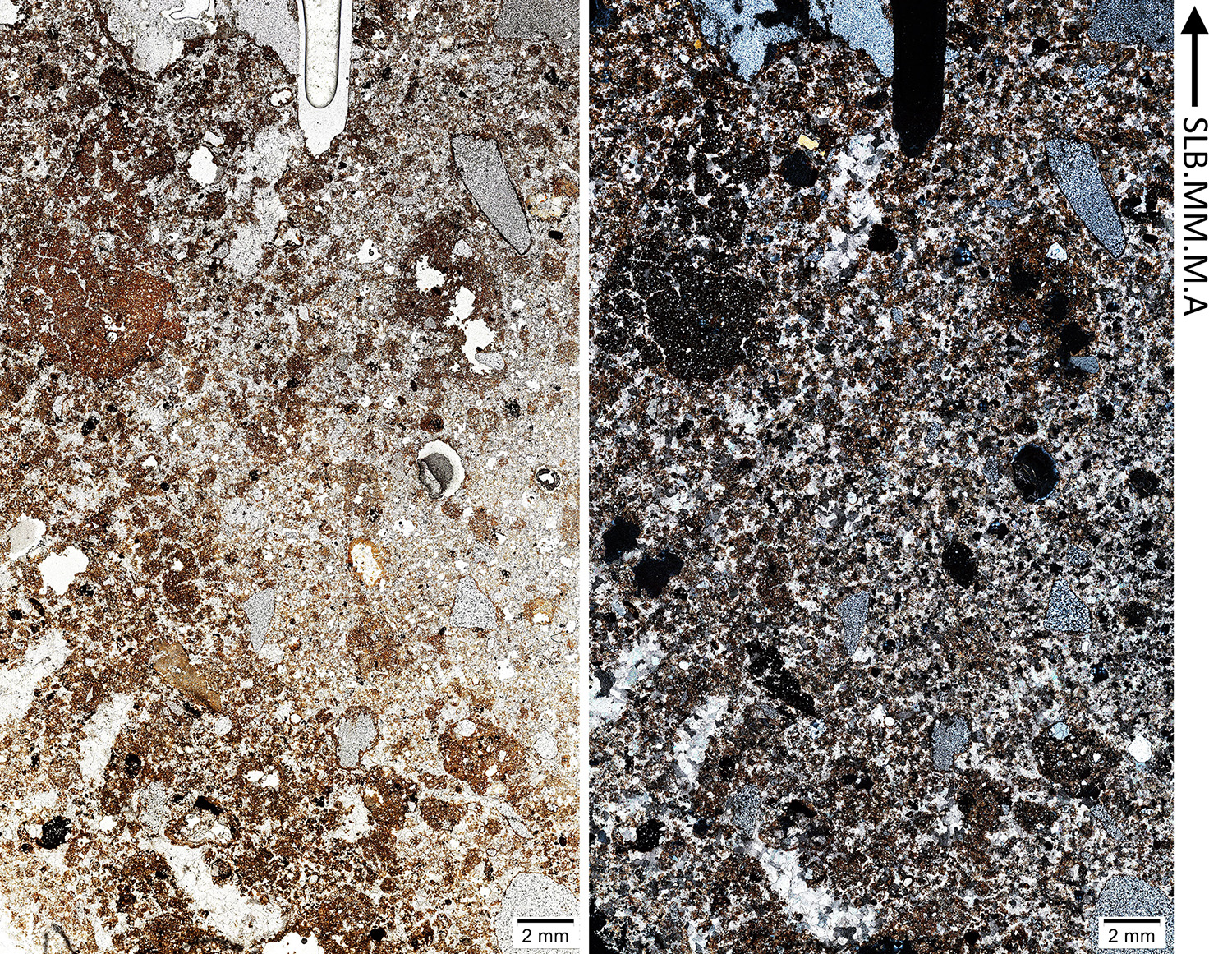

### Supplementary file 4

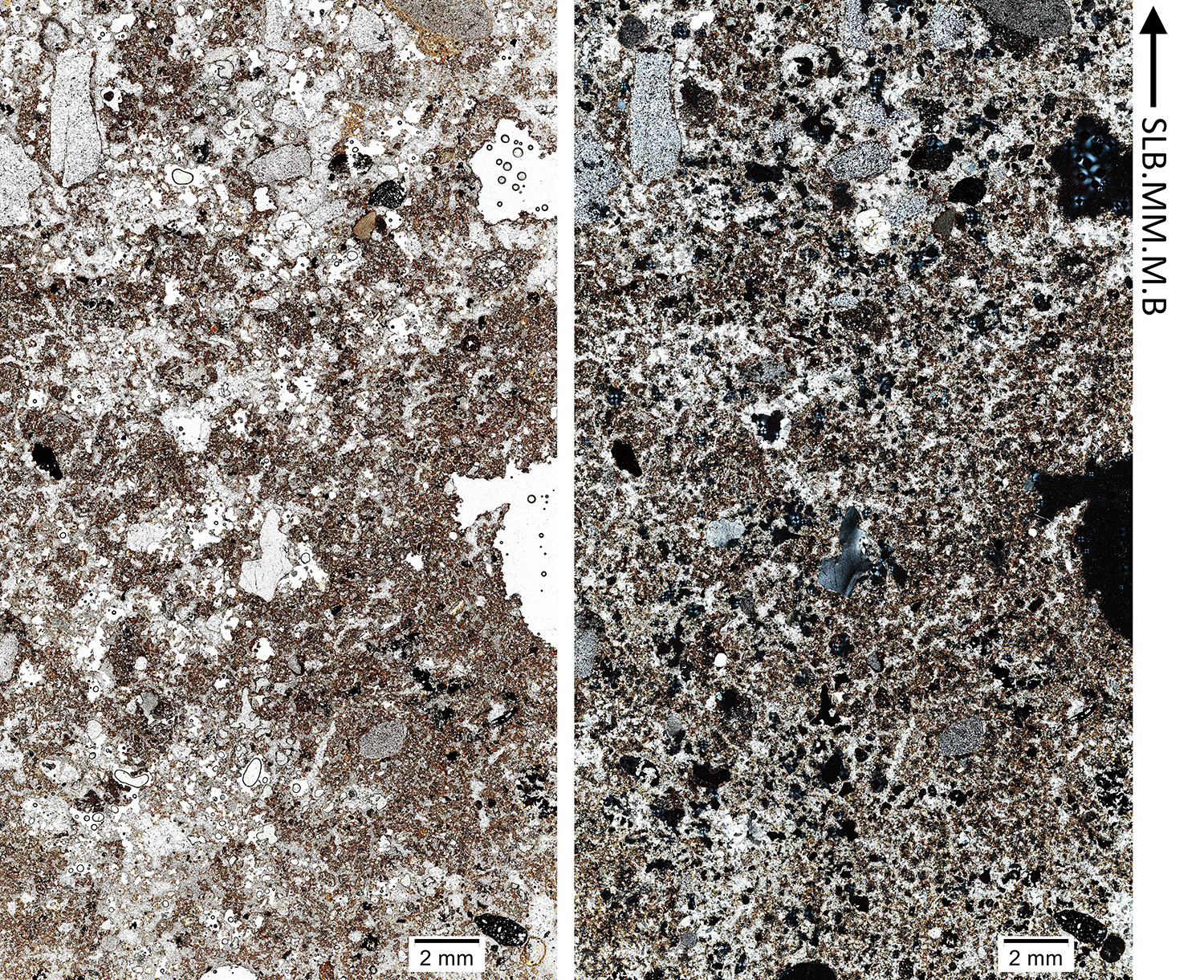

### Supplementary file 5

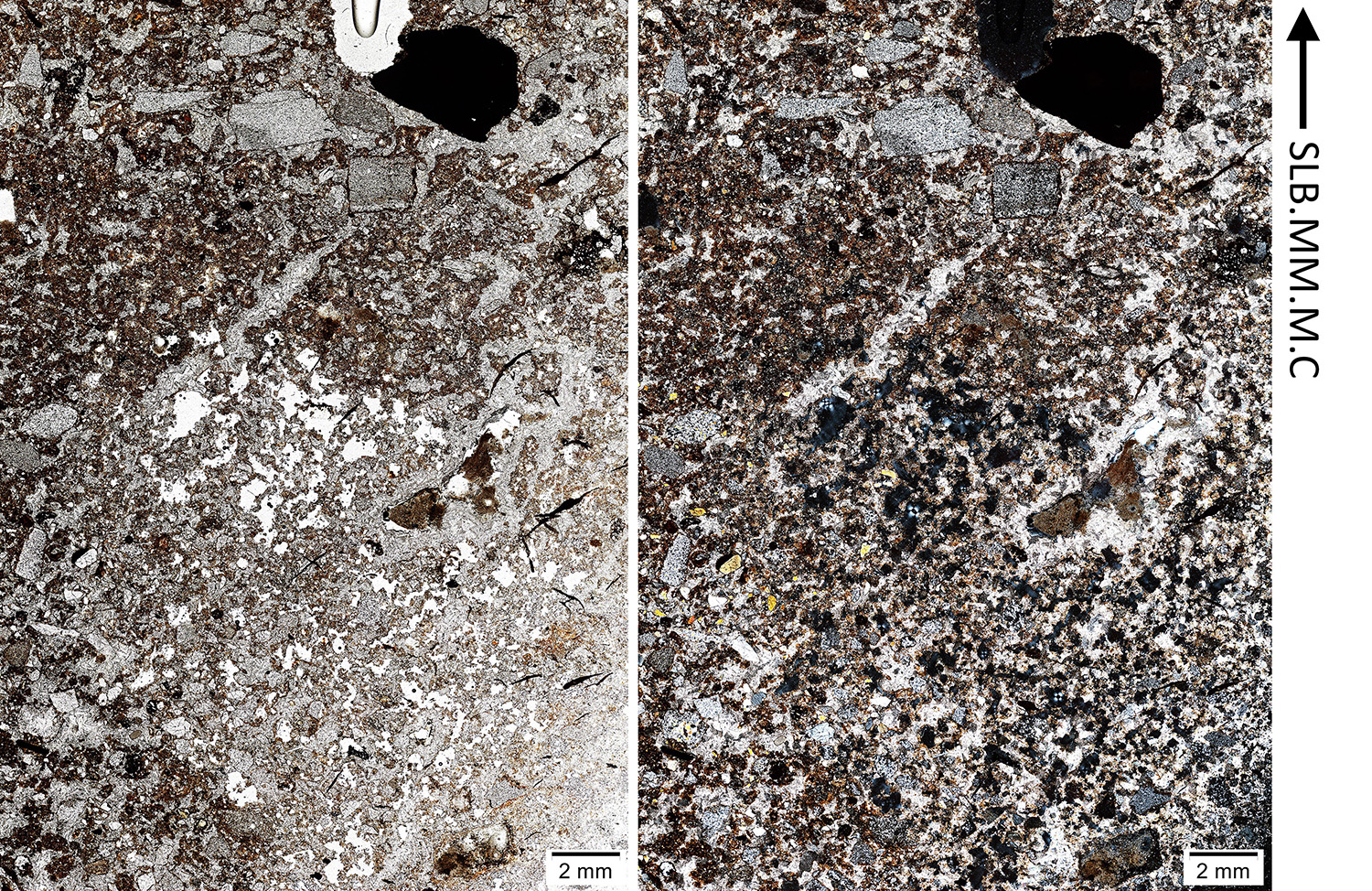
